# Calcium and Bicarbonate Signaling Pathways have Pivotal, Resonating Roles in Matching ATP Production to Demand

**DOI:** 10.1101/2022.10.31.514581

**Authors:** Maura Greiser, Mariusz Karbowski, Aaron D. Kaplan, Andrew K. Coleman, Carmen A. Mannella, W. J. Lederer, Liron Boyman

## Abstract

Mitochondrial ATP production in cardiac ventricular myocytes must be continually adjusted to rapidly replenish the ATP consumed by the working heart. Two systems are known to be critical in this regulation: mitochondrial matrix Ca^2+^ ([Ca^2+^]_m_) and blood flow that is tuned by local ventricular myocyte metabolic signaling. However, these two regulatory systems do not account for the large physiological range of ATP consumption observed. We report here on the identity, location, and signaling cascade of a controversial third regulatory system -- CO_2_/bicarbonate. CO_2_ is generated in the mitochondrial matrix as a metabolic waste product produced by oxidation of nutrients which power the production of ATP. It is a lipid soluble gas that equilibrates with bicarbonate (HCO_3_−) in aqueous solutions. The bicarbonate level is tracked by a bicarbonate-activated adenylyl cyclase, soluble adenylyl cyclase (sAC). Using structural Airyscan super-resolution imaging and functional measurements we find that sAC is primarily inside the mitochondria of ventricular myocytes where it generates cAMP when activated by HCO_3_−. This cAMP signaling cascade is shown to operate inside the mitochondrial inter-membrane space (IMS) by activating local EPAC1 (**E**xchange **P**rotein directly **A**ctivated by **c**AMP) which turns on Rap1 (Ras-related protein 1). Thus, mitochondrial ATP production is shown to be increased by bicarbonate-triggered sAC signaling through Rap1. Additional evidence is presented indicating that the cAMP signaling itself does not occur directly in the matrix. We also show that this third signaling process involving bicarbonate and sAC activates the cardiac mitochondrial ATP production machinery by working independently of, yet in conjunction with, [Ca^2+^]_m_-dependent ATP production to meet the energy needs of cellular activity in both health and disease. Thus, the resonant, or complementary effects of bicarbonate and [Ca^2+^]_m_ signaling arms tune mitochondrial ATP production to match the full scale of energy consumption in cardiac ventricular myocytes.

## Introduction

Cardiac ventricular myocytes work non-stop to meet the blood flow needs of the body, with the heart consuming ATP at high but widely variable rates ^1–11^. These everchanging energy needs, and minimal ATP reserves ^12, 13^, demand that ATP production be dynamically managed by tight, robust, and high-gain feedback controls^14, 15^. Recent work has shown how heart rate dependent elevation of cytosolic Ca^2+^ signaling drives the ventricular myocytes to contract more frequently and, in parallel, the resulting elevated mitochondrial matrix Ca^2+^ ([Ca^2+^]_m_) augments ATP production ^1, 16, 17^. The energetic needs of ventricular myocytes are also linked to local blood supply through the newly described “electro-metabolic signaling (EMS)”. Through this mechanism, as [ATP] in ventricular myocytes falls with ATP consumption, small blood vessels dilate to increase local blood flow ^18, 19^. EMS thereby will increase the supply of oxygen and nutrient substrates, and speed up removal of metabolic waste products by increased flow through local end-arterioles and capillaries. Together, EMS and [Ca^2+^]_m_ signaling provide important feedback-controls to increase ATP production in working ventricular myocytes. There is, however, still a huge gap in accounting for the full physiological scale of ATP consumption by ventricular myocytes ^1–11^. There is no characterized feedback signal yet that provides a mechanism for ventricular myocytes to scale-up ATP production by their mitochondria when increased work is being done by the heart at constant [Ca^2+^]_m_. This kind of situation arises routinely when the heart must pump at the same rate but against a greater afterload. This occurs, for example, when blood pressure rises. The investigation presented in this paper seeks to fill this major gap in our understanding.

Here we report on an understudied and poorly understood signaling pathway in cardiac mitochondria in which CO_2_ and soluble adenylyl cyclase (sAC) play pivotal roles. CO_2_ is generated in mitochondria largely as a waste product originating from processing of energy substrates by the Krebs cycle, therefore reflecting the extent of energy metabolism. This CO_2_ is dissolved in the local aqueous fraction and is in dynamic aqueous equilibrium with bicarbonate (HCO_3_−). Importantly, bicarbonate is the signal sensed by sAC that activates it. sAC is distinguished from the more familiar “trans-membrane” adenylyl cyclase (tmAC) in that it is found in solution. However, like tmAC, when sAC is activated, it too generates cAMP as the second messenger ^20, 21^.

The ten-member family of signaling proteins known as adenylyl cyclases (ACs) is exquisitely adaptable and transduces a wide array of biological signals by generating the second messenger cAMP ^22,23^. The family is best known by the nine members that are incorporated in the plasma membranes of almost every cell type via transmembrane (tm) protein domains. While tmACs are broadly regulated by G proteins in response to hormonal stimuli ^24, 25^, they are able to maintain specificity and high signal-to-noise ratio through proximity to their targets docked at A-kinase anchoring proteins (AKAP’s) ^26, 27^. Further signal compartmentalization is gained by nearby “firewalls” of phosphodiesterases (PDEs) that hydrolyze cAMP to AMP ^28–31^. In contrast, there is only one mammalian member of the soluble adenylyl cyclase subset, and it is much less studied or understood. Mammalian sACs are structural homologues of the bacterial sACs and also lack a transmembrane domain. Instead, they are confined inside organelles like mitochondria, centrioles and nuclei ^32–34^. Unlike tmACs, sACs achieve high local signal-to-noise ratio by producing cAMP inside the small subcellular volumes that also contain their targets ^35, 36^, namely Protein Kinase A (PKA) and “Exchange Proteins Activated by cAMP” or EPACs ^37–39^.

A number of recent papers investigated sAC and its activation by bicarbonate seeking to identify what was sensed by sAC, where in the cell the reactions took place and what the purpose of this signaling system was^**34, 40–42**^. This work by Manfredi and colleagues, although widely cited, is controversial^**23, 43–45**^. These publications concluded that the bicarbonate-activated sAC works within the mitochondrial matrix to produce cAMP, which activates protein kinase A (PKA), which in turn increases ATP production. It was also reported that the role of this signaling system is to sense and respond to “nutrient availability”^**34, 40–42**^. Despite seeming straightforward findings, all key mechanistic findings have been disputed by other investigators^**23, 44 43, 45**^. While all investigators agree that elevated bicarbonate leads to an increase in ATP production, they disagree on how this happens. The role of PKA in the matrix is disputed^**23, 44 43, 45**^ as is the matrix localization of native sAC^**43, 45**^. Furthermore, key pharmacological tools used in the original study were later found to cause acute mitochondrial toxicity rather than selectively inhibit sAC^45, 46^. In light of these major points of disagreement, we investigated sAC quantitatively with high spatial resolution to determine where sAC was located, how it worked, where the intermediate signals were generated, and importantly, what its overall role in cellular metabolism may be. Additionally, we decided to use fresh and functional heart tissue as our source of mitochondria because it is one of the most metabolically dynamic tissues. Additionally, we reasoned that by discovering how sAC worked in cardiac mitochondria, we were likely to provide a solid physiological framework to understand how sAC worked in other tissues. In contrast, the original investigations^**34, 40–42**^ of sAC used mitochondria from cells in which mitochondrial ATP production normally operates over a very narrow range ^47–52^ and sAC signaling itself has low signal-to-noise ratio. To augment the signal, transgenic sAC was overexpressed in the model cells thereby possibly obscuring the role of native sAC.

When each element was studied, we found that all of the key conclusions of the initial investigation by Manfredi and colleagues linking sAC and PKA in the matrix to ATP production were disputed by our data and those of other investigators (see Table-1). Additionally, we found that sAC signaling reports on nutrient *<u>consumption</u>* not on nutrient “availability”^**34, 40–42**^. Furthermore, a major finding of the work presented here is that mitochondrial sAC signaling works independently of, yet in conjunction with, mitochondrial Ca^2+^ signaling system. Together they report on the full scale of ATP needs in health and disease. This resonant or complementary signaling between the two systems enables the mitochondria in cardiac ventricular myocytes to produce ATP continuously at levels appropriate for the physiological demands of the heart.

## Results

An investigation is presented that centers on where sAC is located within the ventricular myocyte and how that position affects its function. That investigation requires high resolution imaging and quantitative biochemical investigations to determine where sAC resides and what its product, cAMP, controls. This work then examines how the local potential targets of cAMP, EPAC1 and PKA, may be connected to the dynamic control of ATP production by mitochondria. Under these conditions, sAC signaling is examined in context of Ca^2+^ signaling to examine their interactions. To place the results in a physiological context they are discussed in the setting of the working heart.

### Localization of Soluble Adenylyl Cyclase (sAC) in ventricular myocytes

Immunofluorescence localization of sAC at high resolution was undertaken using the Zeiss Airyscan 880 super-resolution microscope and the R21 monoclonal anti-sAC antibody ^32, 45, 56–58^ validated in sAC knockout mice ^59^. Figure 1A shows simultaneously acquired images of a ventricular myocyte co-labeled for sAC, ATP synthase (complex-V or C_V_ of the ETC) and F-actin. ATP synthase is a component of the inner mitochondrial membrane (IMM) while F-actin filaments are extramitochondrial. Figure 1A (left panel) shows the location of sAC (yellow), with a zoomed-in region in the upper right corner. The sAC labeling clearly localizes to the mitochondria, which appear with punctate features (roughly 1 μm across) occurring in rows between the F-actin-containing (blue) myofilaments (see Figure 1A) ^60^. This conclusion is strengthened by the similarity of the distribution of sAC to that of ATP-synthase (red in Figure 1A). This point is well-illustrated in the merged image. The mitochondrial localization of sAC is also supported by colocalization analysis measured by the Pearson’s correlation coefficient between C_V_ and sAC (Figure 1B). Likewise, in Figures 1C and 1D it is clear that neither protein is colocalized with the contractile filament (F-actin). This examination of the transverse axis of the ventricular myocytes shows that sAC and C_V_ occur at the same location (Figures 1C) and same frequency in the myocyte. While F-actin is also found at the same frequency as sAC and C_V_, F-actin is only found between them (Figures 1D and 1E). While sAC and ATP synthase are co-localizing to mitochondria, the question remains, where in the mitochondria is sAC located. There are four possible locations to consider: the outer mitochondrial membrane (OMM), the inter-membrane space (IMS), the inner mitochondrial membrane (IMM), and the matrix.

**Figure 1.**
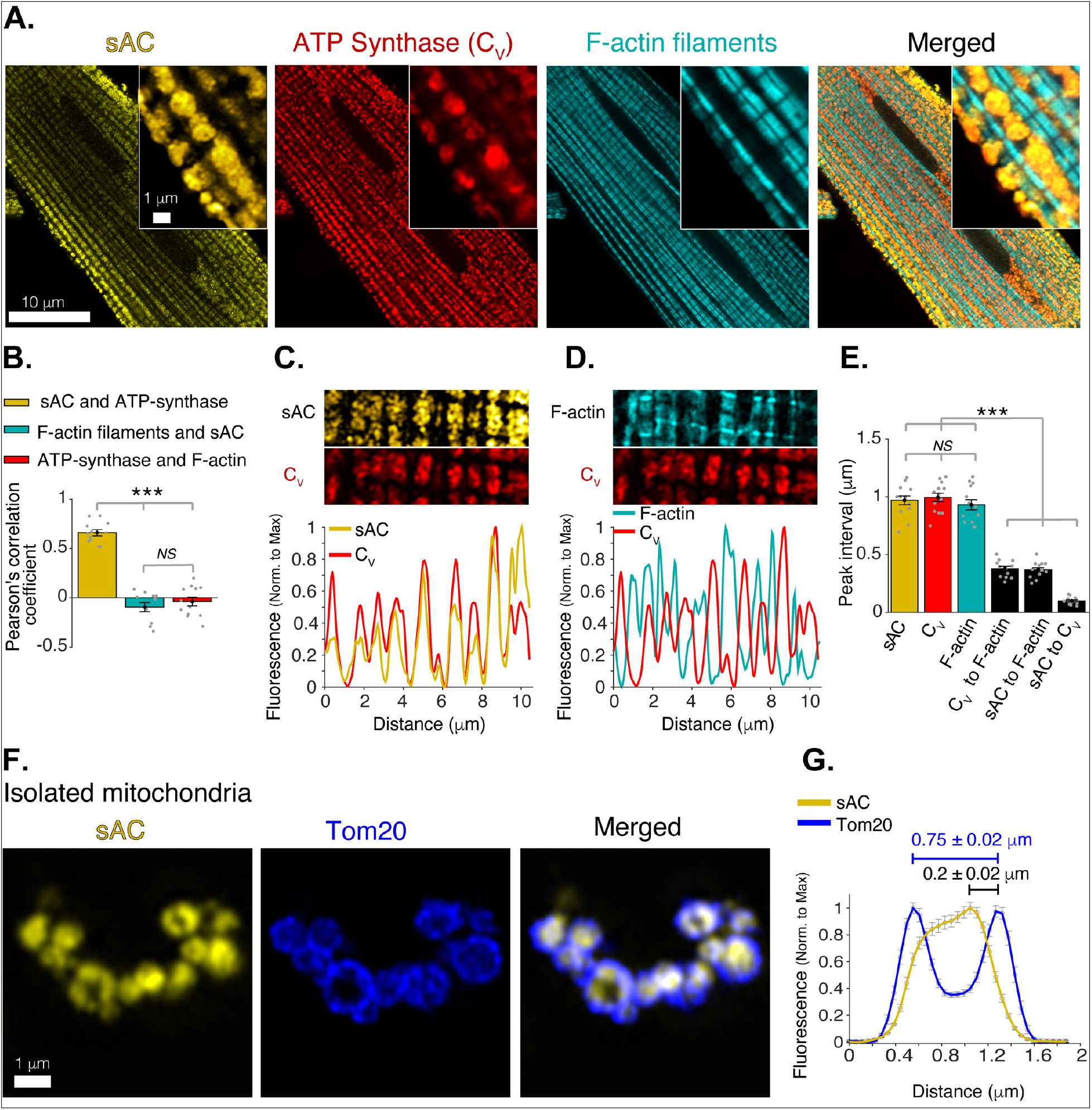
Mitochondrial localization of soluble adenylyl cyclase (sAC). **A**. AiryScan super-resolution fluorescence images of a cardiomyocyte (left to right) immuno-labeled for sAC (yellow) and ATP synthase (red), and loaded with Alexa Fluor™ 488 phalloidin (1 μM) to label the F-actin within the contractile filaments (cyan). Merged image (far right) shows superimposed labeling of sAC, ATP synthase, and F-actin. **B**. Pearson’s correlation analysis of sub-cellular colocalization of sAC with ATP synthase (yellow), sAC with F-actin filaments (cyan), and ATP synthase with F-actin filaments (red), (*n*= 12 cells). **C**. Fluorescence profile of sAC (yellow) and ATP synthase (red). **D**. Fluorescence profile of F-actin (cyan) and ATP synthase (red). **E**. Mean of peak interval for data as in C-D (*n*= 13 cells). **F**. AiryScan super-resolution images of isolated cardiac mitochondria immuno-labeled for sAC (yellow, left), Tom20 (blue, center), and merged images (right). **G**. Fluorescence profile of sAC (yellow) and Tom20 (blue) for individual mitochondrial images as in F, (*n*= 36). Mean peak intervals (in μm) are indicated (*n*= 36 mitochondria). Data in **B, E**, and **G-J** are mean ± SEM. One-way two-tailed ANOVA with Bonferroni correction in **B, E**, and **G-J**. *** P<0.001. NS = not significant (P>0.05).

To narrow down the possibilities, super-resolution microscopy was applied to isolated mitochondria (Figures 1F, 1G), using immunolabeling for sAC and Tom20 (Translocase of the Outer Membrane 20), an abundant OMM protein ^61^. The resolution achieved clearly distinguished between OMMs of adjacent mitochondria and showed that sAC is located within the perimeter of the OMM but is not co-localized with Tom20. Since sAC does not contain any membrane-spanning sequences, it is unlikely to be embedded in the IMM. This then suggests that sAC is in the intermembrane space or in the matrix or in both compartments.

### Bicarbonate Sensing in Isolated Cardiac Mitochondria

There are two important functional distinctions between sACs and tmACs. First, sAC is directly activated by HCO_3_− to produce cAMP while the tmACs are not ^20, 21^. Second, sAC is not activated by forskolin to produce cAMP ^20, 21^, while all 9 types of tmACs are ^62^. We used these established features of the different kinds of ACs to determine the identity of the ACs in cardiac mitochondria. As indicated by the experiments of Figure 2A, treating isolated mitochondria with bicarbonate robustly activates cAMP production (over the physiologic range of bicarbonate from 0 to 15 mM) while treatment with forskolin (25 uM) had no effect on cAMP production. The simple conclusion from these experiments is that the cardiac mitochondria contain functional sAC and no detectable tmACs.

**Figure 2.**
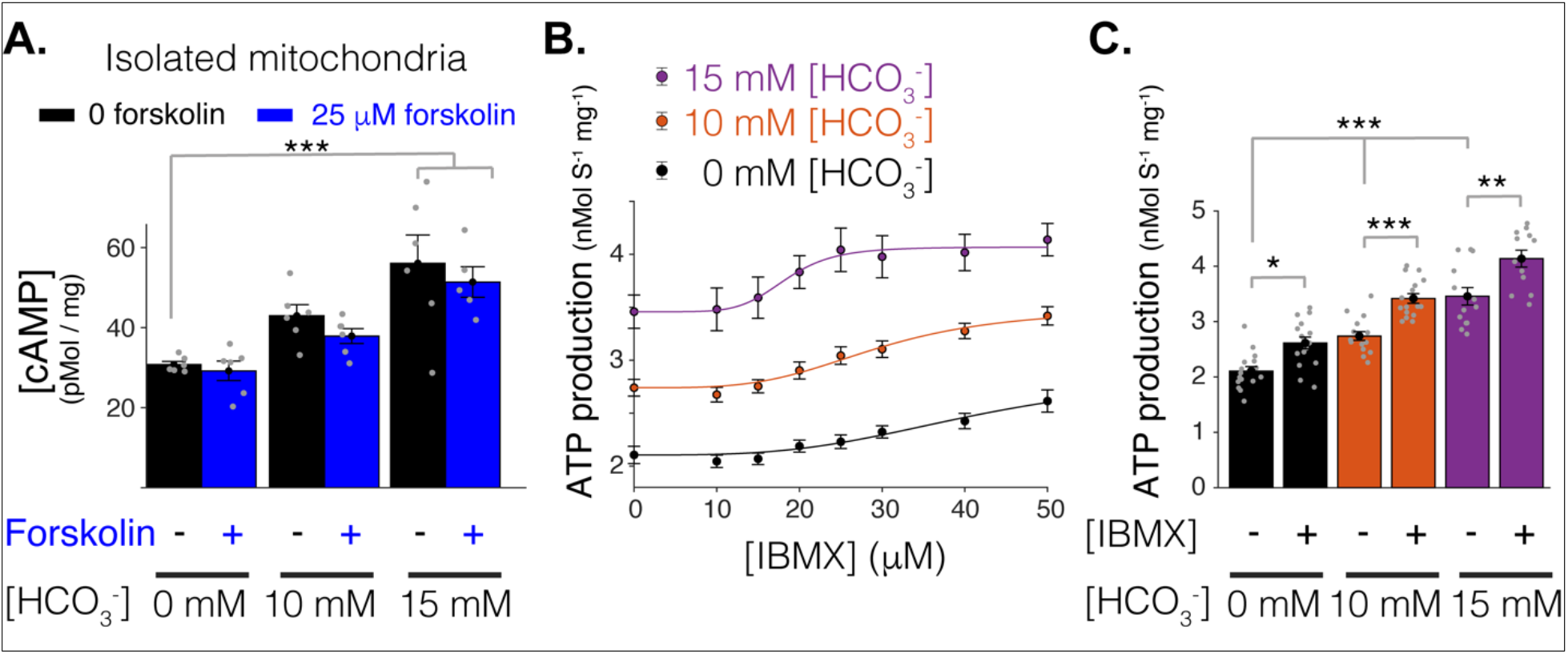
Mitochondrial function of soluble adenylyl cyclase (sAC). **A**. Quantitative ELISA measurements of cAMP inside isolated mitochondria (pMol/mg). cAMP is measured following treatment with indicated concentrations of bicarbonate (HCO_3_−), which activates soluble adenylyl cyclase (sAC), and forskolin, which activates transmembrane adenylyl cyclases (tmACs). (*n*= 6, 6, 5 for 0, 10, 15 mM [HCO_3_−], respectively, mitochondria are isolated independently from 4 hearts). **B**. The dependence of ATP production on [HCO_3_−] and [IBMX]. Sigmoidal fits to the data are shown. **C**. The sensitivity of ATP production to phosphodiesterase inhibition (with 50 μM IBMX) at different concentrations of [HCO_3_−]. For **B-C***n*= 12-16 independent experiments per group, mitochondria are isolated independently from 4 hearts. Data in **A-C** are mean ± SEM. One-way two-tailed ANOVA with Bonferroni correction in **A** and **C**. * P < 0.05, ** P<0.01, *** P<0.001.

Further definition of the behavior of the mitochondrial sAC system was provided by use of the membrane permeable phosphodiesterase (PDE) inhibitor IBMX, 3-Isobutyl-1-methylxanthine ^63^. PDEs are plentiful in nearly all cells that use cAMP and are distributed to help focus the action of the cyclic nucleotide to a local region ^29, 64^. The PDEs limit the diffusion of the cAMP away from their intended target. This has often been likened to a “firewall” against the excessive signaling of cAMP within a region of a cell or within an organelle ^28–31^. Conversely, inhibition of endogenous PDEs within mitochondrial compartments would be expected to enhance cAMP signaling within those compartments. The effects of the PDE inhibitor IBMX on the action of sAC in isolated mitochondria are presented in Figures 2B and 2C. These results provide primary evidence that ATP production by cardiac mitochondria is enhanced by bicarbonate-activated sAC and regulated by PDEs. Specifically, physiological concentrations of HCO_3_− (10 and 15 mM) significantly increase ATP production, and IBMX elevates it further, for a combined effect of doubling ATP output. These data are consistent with a cAMP signaling system inside mitochondria that responds to bicarbonate by activating sAC, that is modulated by PDEs that reduce cAMP levels, and that regulates the primary function of these mitochondria, the generation of ATP. The response of mitochondria to sAC signaling is such that, as [HCO_3_−] increases, so too will ATP production. In this manner sAC can track CO_2_ production (as [HCO_3_−] levels) and thereby upregulate mitochondrial ATP production in response to increased energy consumption (substrate oxidation) and thus metabolic need.

### Mitochondrial cAMP signaling

The above results indicate that the cAMP generated inside mitochondria by sAC and regulated by PDEs directly modulates ATP production by mitochondria. To investigate this process in more detail, the characteristics of the sAC signaling system aid us in the design of experiments to quantitatively assess the contributions of sAC to metabolic regulation of the heart. A critical characteristic is that the sAC output signal following activation is cAMP, which cannot readily cross a membrane due to its hydrophilicity ^44, 46^. We can use this information to determine in which compartment sAC signaling is working in ventricular myocytes. In the special case of an isolated mitochondrial preparation, nucleotides like cAMP applied to the extra-mitochondrial space gain entry to the mitochondrial inter-membrane space (IMS) through the numerous VDAC pores in the outer mitochondrial membrane (OMM) ^53–55^. However, cAMP has been shown to be excluded from the matrix because the inner mitochondrial membrane (IMM) is impermeable to it ^44, 46^. The experiments shown in Figure 3 use this understanding to investigate sAC signaling in more detail.

**Figure 3.**
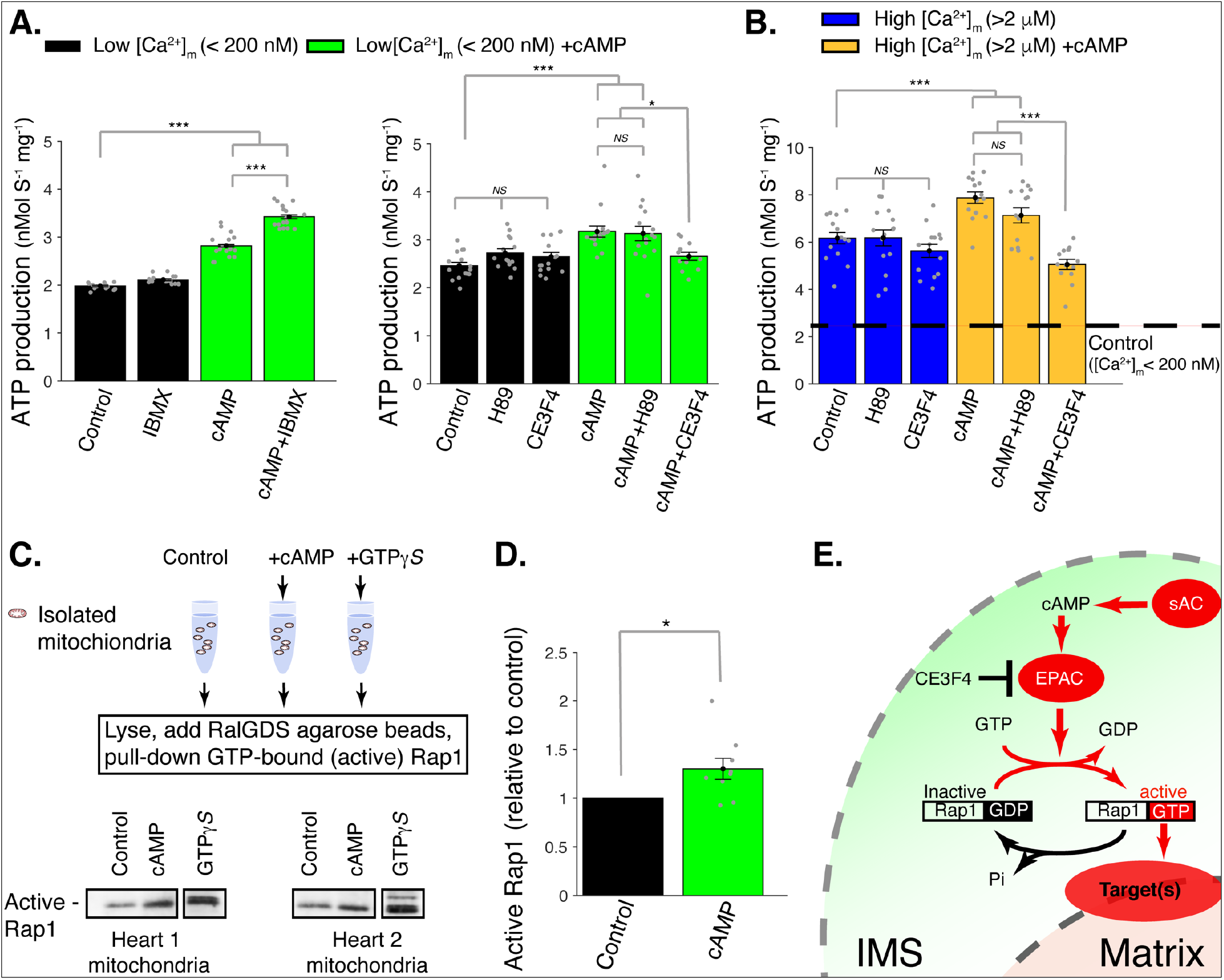
cAMP control of mitochondrial ATP production. **A**. Left, sensitivity of mitochondrial ATP production to cAMP and phosphodiesterase inhibition (with 50 μM IBMX). Measurements are carried out at low [Ca^2+^]_m_ (< 200 nM).Right, sensitivity of mitochondrial ATP production to cAMP, EPAC1 inhibition (25 μM CE3F4), and PKA inhibition (1 μM H89). Measurements were carried out at low [Ca^2+^]_m_ (< 200 nM). **B**. Same as A but at high [Ca^2+^]_m_ (> 2 μM). **C**. Pull-down and immunoblot analysis for the active form of Rap1 (GTP-bound Rap1) in isolated mitochondria stimulated with cAMP. γ-GTP is used to maximally activate Rap1. **D**. The relative amounts of active Rap1 (mitochondria isolated independently from n = 9 hearts). **E**. Schematic diagram showing the likely locations of key proteins in sAC mitochondrial signaling. While sAC is clearly located within the intermembrane space (IMS), and both EPAC and Rap1 have signaling domains in the IMS, it is uncertain how the “target” protein(s) are activated to increase ATP production. For **A-B***n*= 12-16 independent experiments per group, mitochondria are isolated independently from 4 hearts. In **D** Mitochondria are isolated independently from 9 hearts. Data in **A-B**, and **D** are mean ± SEM. One-way two-tailed ANOVA with Bonferroni correction in **A-B**. One-Sample t-Test for **D**. * P < 0.05, ** P<0.01, *** P<0.001. **Figure 3—source data 1. Western blot analysis in Figure 3C-D (anti-Rap1).** **Figure 3—source data 2. Original file No.1 for western blot analysis in Figure 3 C-D (anti-Rap1).** **Figure 3—source data 3. Original file No.2 for western blot analysis in Figure 3 C-D (anti-Rap1).** **Figure 3—source data 4. Original file No.3 for western blot analysis in Figure 3 C-D (anti-Rap1).** **Figure 3—source data 5. Original file No.4 for western blot analysis in Figure 3 C-D (anti-Rap1).**

Figure 3A, shows that ATP production in isolated mitochondria with low [Ca^2+^]_m_ is small, about 2 nM ATP mg^−1^ s^−1^ and is not significantly increased by the PDE inhibitor IBMX. These data suggest that there is little or no cAMP present in the mitochondrial compartment in which sAC is located under these conditions. However, when cAMP is added extra-mitochondrially to volume that also contains the mitochondria, there is a clear increase of approximately 50% in ATP production that is further augmented to 100% (doubling) upon addition of IBMX. This finding re-emphasizes that the sAC is in the mitochondria and, moreover, that it is located in the IMS, i.e., accessible to externally added cAMP. While nucleotides can enter the IMS through VDAC ^53–55^, it has been shown that cAMP cannot cross the IMM ^44, 46^. Thus, the production of cAMP by sACs activates a target in the IMS which, in turn, produces a signaling cascade that can activate one or more targets in the IMM and/or matrix space.

There are two possible direct targets for this locally elevated cAMP -- namely, PKA (Protein Kinase A) and/or EPAC (**E**xchange **P**rotein directly **A**ctivated by **c**AMP). To examine these targets, experiments were conducted under conditions where matrix Ca^2+^ levels ([Ca^2+^]_m_) were measured quantitatively and kept low (under 200 nM). Figure 3A (right panel - black bars) shows that, in the <u>*absence* of added extra-mitochondrial cAMP</u>, blocking PKA (with its inhibitor H89) does not affect ATP production nor does the blocking of EPAC1 by the inhibitor CE3F4. However, when cAMP is applied extra-mitochondrially in low [Ca^2+^]_m_ (green bars) there is an increase in ATP production that is inhibited by CE3F4 but not H89. From these experiments we conclude that mitochondrial EPAC1 is the target protein activated by cAMP applied to isolated mitochondria, and that PKA is not involved. We also conclude that EPAC1 contributes to increased generation of ATP even in low [Ca^2+^]_m_ and that this signaling happens within the IMS. We conclude that the signaling is in the IMS because cAMP does not enter the matrix under the conditions of our experiment.

Figure 3B shows that at elevated [Ca^2+^]_m_ (> 2 μM), in the absence of cAMP (blue bars), there is a significant increase (2.6-fold) in ATP production compared to the low [Ca^2+^]_m_ control (dashed black line). There is an important further increase in ATP production (1.3-fold - brown bars) when cAMP is added to these mitochondria with elevated [Ca^2+^]_m_. These findings suggest that IMS sAC signaling and the [Ca^2+^]_m_ signaling system have additive (independent) effects on ATP production. This conclusion is also supported by our findings that the sAC signaling system works without the need for elevated [Ca^2+^]_m_ (Figures 2–3). In addition, as shown in Figure 3A and 3B, the cAMP-dependent increase in ATP production was sensitive to EPAC1 inhibition. The pull-down and subsequent immunoblot data in Figures 3C and 3D show that this action can be attributed to activation of the canonical EPAC1 target protein, a member of the Ras-related protein family, Rap1 (Repressor/activator protein 1). Diagrammatically this activation process is shown in Figure 3E. While we have shown that much of the signaling (sAC, EPAC1 and Rap1) occurs in the IMS, the exact location and identity of the Rap1-target protein(s) remains unknown and will be examined in future studies.

### cAMP signaling domains: IMS versus matrix

Given our immunolocalization results in Figure 1 using both isolated ventricular myocytes and isolated mitochondria and the functional experiments on isolated mitochondria in Figures 2–3, the sAC -> EPAC1 -> Rap1 signaling pathway appears to be in the IMS. In contrast to our simple results, other studies have suggested significantly more complex results that are in conflict with our findings. For example, some experiments suggest that cAMP signaling includes both cytosolic and mitochondrial matrix targets ^34, 42, 44, 46, 65–67^. To sort out possible contributions from matrix-localized cAMP signaling, we employed the membrane permeant analog of cAMP, 8Br-cAMP. When applied to an isolated mitochondrial preparation, 8Br-cAMP would be expected to activate both EPAC and PKA in all compartments (i.e., both IMS and matrix) of the mitochondria. 8Br-cAMP also is a persistent activator of EPAC and PKA since it is resistant to breakdown by PDEs ^68–70^. Figure 4B shows the effects of 8Br-cAMP application to isolated mitochondria in low [Ca^2+^]_m_ (< 200 nM). In contrast to the application of cAMP extra-mitochondrially, which produces significant increase in ATP production (Figure 3), 8Br-cAMP produces a modest reduction in ATP production. Further, more significant reduction in ATP production is seen with 8Br-cAMP and addition of the PKA inhibitor H89 or the EPAC inhibitor CE3F4 (Figure 4B, left panel). When the PDE inhibitor IBMX is applied (Figure 4B, right panel), it modestly increases ATP production when [Ca^2+^]_m_ is low in the absence or presence of 8Br-cAMP, suggesting persistent effects of cAMP produced in the IMS by sAC and undegraded by PDEs under these conditions. The progressive decrease in ATP production following addition of 8Br-cAMP alone and with blockers of PKA (H89) or EPAC (CE3F4) is also observed at the elevated ATP production rates associated with high [Ca^2+^]_m_ (>2 μM), as shown in Figure 4C. The simple conclusion is that the action of cAMP in the IMS is overwhelmed by the action of 8Br-cAMP in the matrix. Exactly what 8Br-cAMP does in the matrix to produce its effect is not yet known. It is evident, however, that the actions of bicarbonate on sAC, cAMP on EPAC1 and EPAC1 on Rap1 all take place in the IMS. Taken together, the physiological actions of bicarbonate and externally added cAMP that stimulate ATP production by cardiac mitochondria appear to be due to IMS-based signaling, and not to previously described actions of cAMP within the matrix ^34^.

**Figure 4.**
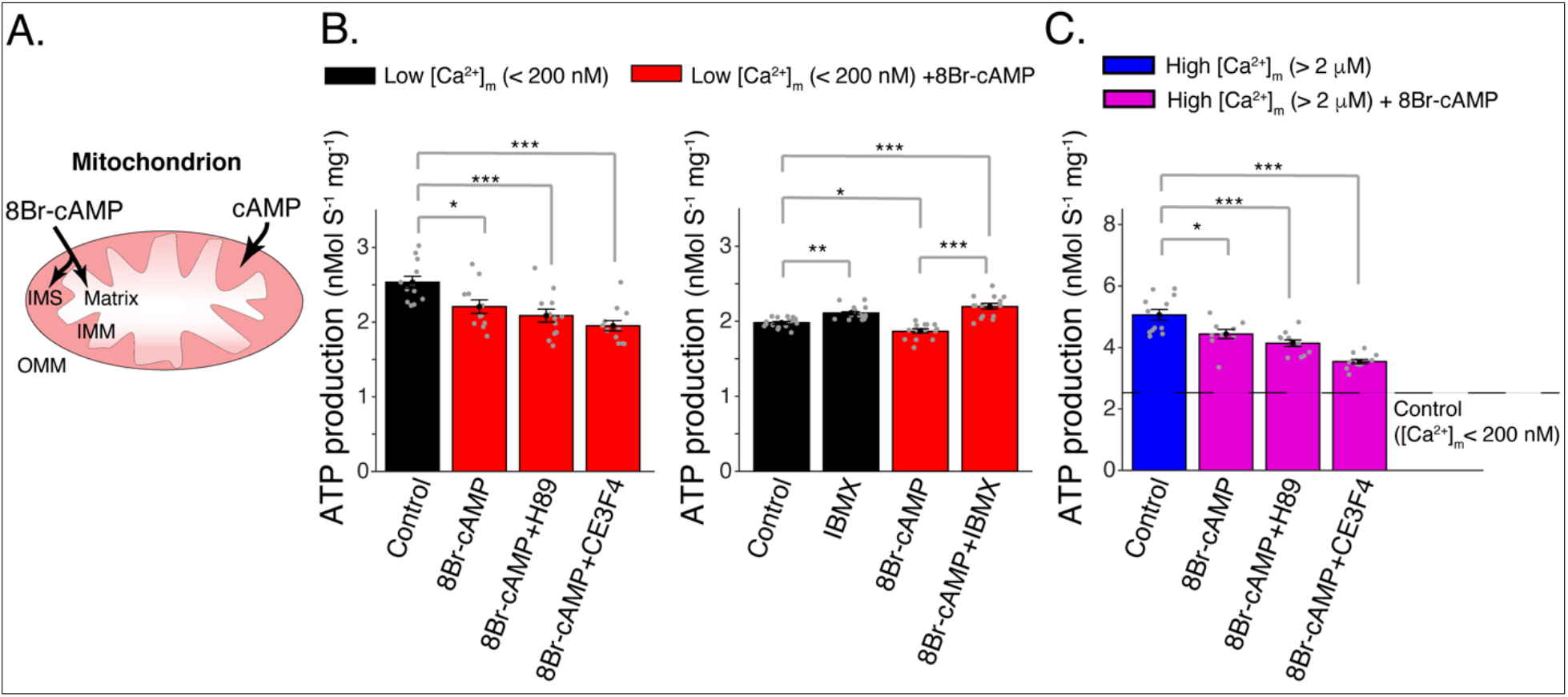
Elevated matrix [cAMP] inhibits mitochondrial ATP production. **A**. Diagram showing mitochondrial compartments accessible to cAMP and the membrane permeant analog, 8Br-cAMP. **B**. Left, mitochondrial ATP production was decreased by application of. This effect was not changed by either EPAC1 inhibition (25 μM CE3F4) or PKA inhibition (1 μM H89). Measurements were carried out at low [Ca^2+^]_m_ (< 200 nM). Right, application of phosphodiesterase inhibitor (50 μM IBMX) produced a small increase in ATP production. Measurements were carried out at low [Ca^2+^]_m_ (< 200 nM). **C**. Same as **B** (left panel) but at high [Ca^2+^]_m_ (> 2 μM). For **B-C***n*= 12-16 independent experiments per group, mitochondria are isolated independently from 4 hearts. Data in **B-C** are mean ± SEM. One-way two-tailed ANOVA with Bonferroni correction in **A-C**. * P < 0.05, ** P<0.01, *** P<0.001.

## Discussion

This investigation has addressed a long-standing metabolic conundrum that is particularly vexing for heart function: how does the consumption of energy contribute to the feedback control of ATP production by mitochondria? The Ca^2+^ signaling pathway is clearly important and depends on Ca^2+^ entry into the mitochondrial matrix to regulate ATP production^1, 16, 17, 46, 71–74^. Here we describe a substrate-consumption-dependent signaling mechanism that provides complementary information to the mitochondria and broadens their range of ATP production. We found that cAMP signaling by soluble adenylyl cyclase (sAC) located in the intermembrane space (IMS) of mitochondria appears to set a scale-factor for ATP generation that increases directly with substrate consumption. Soluble AC has two features that make it an ideal metabolic sensor to complement the [Ca^2+^]_m_ signal for regulation of mitochondrial ATP production under normal conditions. The first is sAC’s distinctive activation by the metabolic waste product CO_2_/HCO_3_−. In effect, bicarbonate sensitivity enables sAC in the IMS to “monitor” the carbon emissions of the adjacent ATP-producing powerplant. Through this mechanism sAC senses an integrated energy consumption “index” in its subcellular environment that reflects the level of cellular CO_2_/HCO_3_− production (as mitigated by vascular waste product removal). Secondly, sAC generates a powerful, locally acting second messenger, cAMP, as an output signal, based on the HCO_3_− that it senses. Importantly, cAMP as a hydrophilic signaling molecule cannot readily cross the inner mitochondrial membrane (IMM) to directly regulate matrix proteins ^44, 46^. Instead, sAC operates with high signal-to-noise ratio over the nanometer scale of the IMS to activate EPAC1 which, in turn regulates guanine-nucleotide exchange factors for the Ras-like GTPase, Rap1. Since neither EPAC1 nor Rap1 has transmembrane domains, the working hypothesis is that Rap1 has an as yet unidentified target in the inner mitochondrial membrane (IMM) that affects ATP synthesis either directly or indirectly, by transmitting its signal to another unknown target in the matrix. The additional significance of these results is that sAC appears to affect mitochondrial ATP production in a manner that does not depend on [Ca^2+^]_m_ but instead complements it. In effect, there are two signaling arms for feedback regulation of cardiac ATP production under normal conditions: one that depends on changes in [Ca^2+^]_m_ (associated with heart rate) and another that depends on variations in CO_2_/HCO_3_− levels (energy consumption), presented schematically in Fig. 5. Thus, our findings provide an important new framework for understanding the dynamic regulation of ATP production in heart and perhaps other tissues. At the same time, this work presents obvious challenges in terms of understanding how the disparate signals are integrated to precisely regulate mitochondrial ATP production over its full dynamic range.

**Figure 5.**
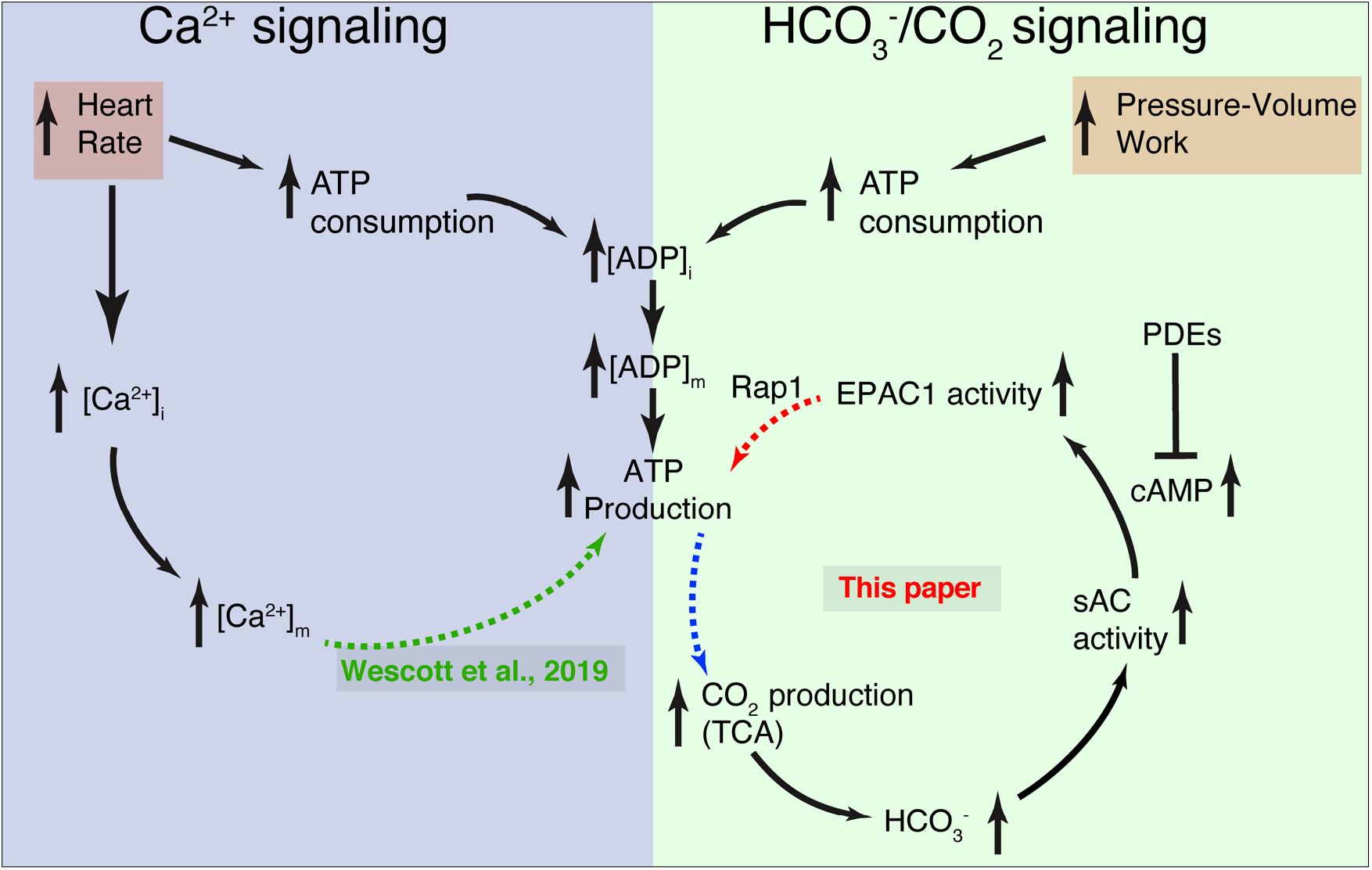
Dynamic control of mitochondrial ATP production by Ca^2+^ and bicarbonate. Schematic diagram of the two complementary signaling arms that control mitochondrial ATP production. **Left**: The Ca^2+^ arm depends on heart rate. With higher heart rates, increased mitochondrial matrix Ca^2+^ ([Ca^2+^]_m_) augments ATP production via a mechanism described in detail in Wescott et. al., 2019. **Right**: The bicarbonate arm responds to pressure/volume work, highlighting the pivotal role played by CO_2_ (in dynamic equilibrium with bicarbonate). This signaling arm is centered in the IMS (intermembrane space between the IMM and the OMM) with the final link (involving Rap1) reaching across the IMM and probably into the matrix. Neither the Ca^2+^ nor the HCO^3−^ signaling arm alone accounts for the full scale of enhancement of ATP production during normal physiological function. Each arm can be stimulated by distinct and different signals, keeping ATP production in sync with heart rate and pressure-volume work. Importantly, it is the resonance signaling between the two signaling arms that properly matches ATP production to actual energy consumption.

### [Ca^2+^]_m_ and CO_2_/HCO_3_− regulatory mechanisms work together: resonance

Under physiological conditions in heart, increasing workload and heart rate can increase mitochondrial ATP production through [Ca^2+^]_m_ -dependent signaling ^1, 16^. In Figure 5, this is represented by the dashed green arrow (bottom, left panel). The elevated [Ca^2+^]_m_ acts on pyruvate dehydrogenase and glutamate dehydrogenase to increase NADH production and thereby increase proton extrusion through the ETC to hyperpolarize the IMM potential, ΔΨ_M_. As ΔΨ_M_ becomes more hyperpolarized (i.e. more negative), ATP production by the ATP synthase increases ^1, 16, 17^. In a separate signaling process, increased pressure-volume work increases ATP expenditure, elevating ADP levels and substrate oxidation rate. Increased CO_2_/HCO_3_− levels activate sAC which generates cAMP and starts a signaling cascade that further increases ATP production by mitochondria (Fig. 5, circular pathway, right panel). This signaling CO_2_/HCO_3_− is produced by the reactions of the TCA (tricarboxylic or Krebs cycle) and pyruvate decarboxylation, both in the mitochondria, as these processes break down and process higher energy substrates. We found that the “bicarbonate” signaling arm of ATP production has the capacity to double the amount of ATP generated per unit time compared to elevated [Ca^2+^]_m_ alone. When considering the two systems working together, there are likely to be many time-dependent and activity-dependent interactions that favor the influence of the [Ca^2+^]_m_ arm or the bicarbonate arm of the ATP production machinery. This gives rise to the idea of resonance signaling between the two arms. The immediate and long-term adaptations of the systems are likely to play an important role in health and disease, as well as in physical training and organ-level adaptation of nuclear and mitochondrial gene expression and protein production. Thus, we have broadly presented the notion of the co-existing processes and accompanying cellular and mitochondrial interactions as a kind of “resonance”. We expect the overall response of ATP production to the two signaling arms (in terms of time- and magnitude-dependence) to be complex and perhaps not totally independent. Sorting out the control elements and mechanisms that produce the changes will provide a deeper physiological understanding and stimulate novel therapeutic opportunities.

### Myocardial Infarction and ATP Production Mechanisms

Our findings presented in Figures 1–4 suggest that there is a clear need for the two independent systems to control mitochondrial ATP production -- one that depends on [Ca^2+^]_m_, and another that depends on HCO_3_−. We established these systems by carrying out experiments on ventricular myocytes and mitochondria that were harvested from healthy hearts. However, to broaden our understanding of how these two systems may work together during cardiovascular stress, we also examined mitochondria from diseased hearts. To do this, we measured the functions of both systems in mitochondria taken from “remodeled” heart tissue in undamaged regions following a myocardial infarction (MI). These results were compared to the same heart region taken from sham operated control hearts. This specific aspect of our discussion is intended to probe the metabolic duality presented in Figures 1–5 and raise additional points of interest for us and for other investigators. It is by no means part of a comprehensive investigation into remodeling of the heart following MI.

MI was produced by ligation of the left anterior descending (LAD) artery ^75, 76^ and examination was done eight weeks post-surgery. Figure 6A shows the large infarct in the anterior wall of the left ventricle. Figures 6B-6C show data from sham versus post-MI hearts and reveal a significantly decreased ejection fraction and a significantly decreased fractional shortening of the post-MI left ventricle. Strain analysis is shown in SI Figure 2. Sample tissue used for measurements was taken from the septum and the posterior left ventricle, far from the scar and the MI border zone. The energy demands of these regions were significantly elevated during the 8 weeks following the MI, largely due to compensation for mechanical dysfunction of the scar tissue and the MI border zone. With this approach, our work was directed at testing how mitochondrial ATP production was regulated by [Ca^2+^]_m_ and HCO_3_− signaling systems in myocytes with persistently high energetic demands. In these measurements we found that the mitochondrial ATP production in the absence of cAMP, and at low [Ca^2+^]_m_ (i.e. <200 nM), was elevated above the sham control levels. Central protein components of the ATP production machinery were largely unchanged (SI Figure 3). These findings could reflect a change in either regulatory system produced by the post-MI remodeling. It is also possible that both systems had changed because the ATP production that is regulated by CO_2_/HCO_3_− and that regulated by [Ca^2+^]_m_ are summed at all measurements. In contrast, the ATP production at high [Ca^2+^]_m_ (i.e. >2uM) was elevated to the same levels as control. We also examined the ATP production regulated by CO_2_/HCO_3_− (Figure 6D) and found that it was increased by application of cAMP. Taken together, these findings are in support of the argument that the CO_2_/HCO_3_− system was working as it had been under non-MI conditions. Thus, in all circumstances (i.e. pre-MI and post-MI and sham-MI), both cAMP, as well as [Ca^2+^]_m_, are needed to control ATP production (see discussion below) but internal set-points are different.

**Figure 6.**
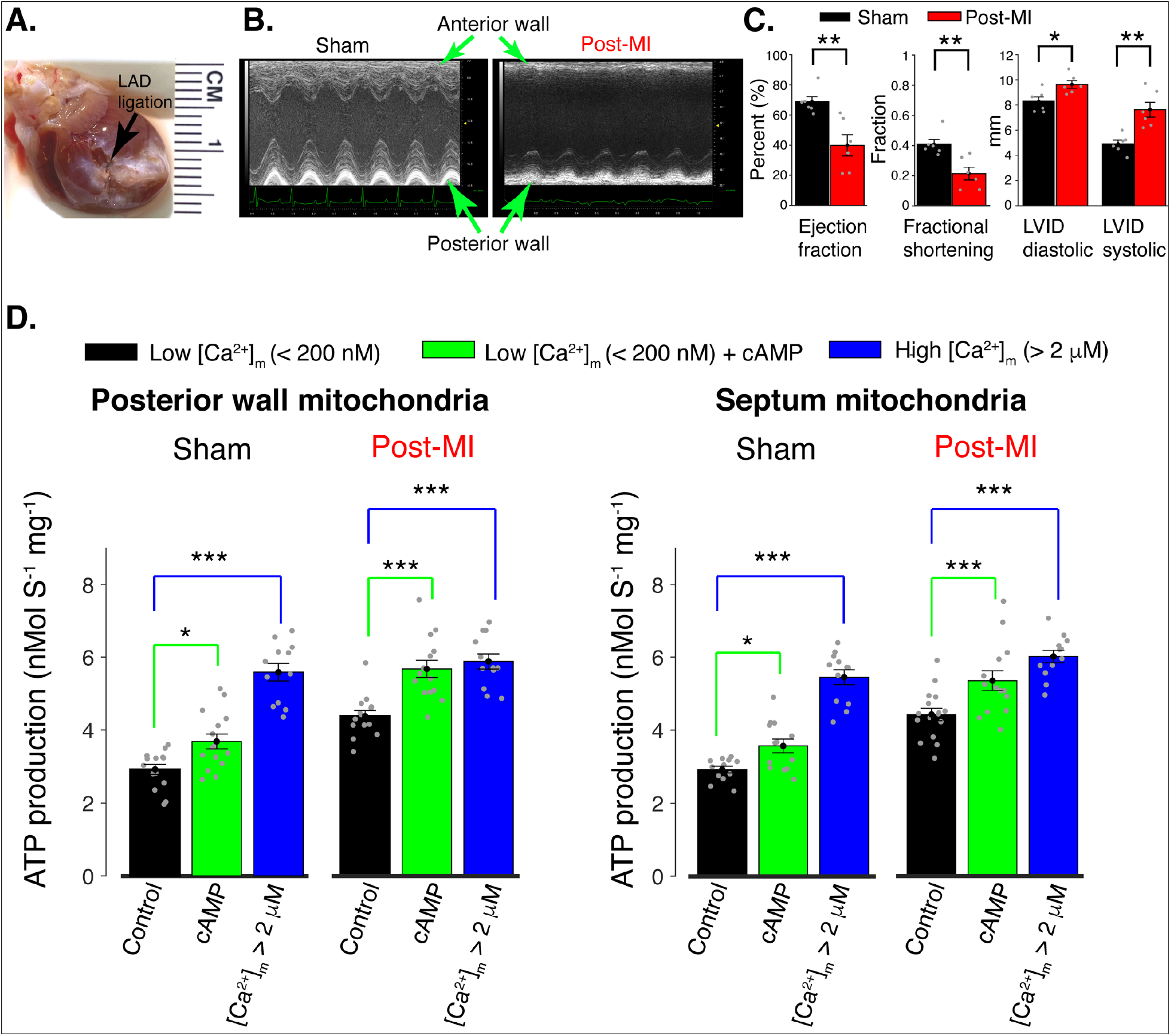
Regulation of mitochondrial ATP production in the post MI heart by [Ca^2+^]_m_ and [cAMP]. **A**. Anterior view of rat heart 8 weeks after induction of myocardial infarction (MI) by ligation of left anterior descending (LAD) coronary artery. The black arrow indicates the location of the ligation and the distal white-colored region shows scar tissue. **B**. Parasternal short-axis M-mode echocardiographic images 8 weeks after surgery involving no ligation (“Sham”) and 8 weeks after genuine LAD ligation (“Post-MI”). **C**. Left ventricular echocardiographic measurements from 6 sham rats (black bars) and from 6 genuine MI rats (red bars). Data are shown as grey circles. Left chart: left ventricle ejection fractions and fractional shortening. Right chart: actual diastolic and systolic left ventricular internal dimensions (LVID). All echocardiographic measurements were carried out under light sedation (1% isoflurane in 100 % oxygen). Heart rate (BPM ± SD): sham 318 ± 28, post-MI 317±12. **D**. ATP production rate in isolated mitochondria at low [Ca^2+^]_m_ (<200 nm, black bars), at low [Ca^2+^]_m_ plus cAMP (green bars), and at high [Ca^2+^]_m_ (>2 μM, blue bars). Mitochondria were isolated from healthy tissue in the septal wall or posterior wall 8 weeks post-MI or post sham operation, as indicated. For all data *n* = 12–16 independent experiments per group, 4 post-MI hearts and 4 sham hearts. Data in **C-D** are mean ± SEM. Two-tailed unpaired t-test in **C**. One-way two-tailed ANOVA with Bonferroni correction in **D**. * P < 0.05, ** P<0.01, *** P<0.001.

### Mitochondrial localization and function of sAC

Given the highly localized nature of cAMP signaling ^26–31^ considerable effort was made to pinpoint the sub-mitochondrial localization of sAC. We were able to determine with super-resolution imaging that sAC is predominantly in the interior and not the outer periphery of mitochondria of ventricular myocytes, suggesting its confinement to the matrix and/or intermembrane space (IMS). A more precise localization was not possible using this approach due to the tight packing of inner membrane compartments (cristae) in cardiac mitochondria, which we could not resolve optically.

Subsequent biochemical results shown here provided strong evidence that sAC proteins were located in the mitochondrial IMS where they generate cAMP that activates EPAC1 in the same compartment. This is the first report of an ATP regulating signaling system in the mitochondrial IMS. In our experiments cAMP applied in the extramitochondrial solution should passively diffuse into the IMS through the large VDAC pores in the outer mitochondrial membrane ^53–55^ but cannot permeate the inner mitochondrial membrane (IMM) ^44, 46^. Conversely, 8Br-cAMP is a cAMP analogue that is broadly used in experiments with intact cells and organelles due to its relatively high membrane permeability ^68–70^. When added to cardiac mitochondria, 8Br-cAMP presumably enters both IMS and matrix compartments, and as we found, causes ATP production to decline. Clearly, its ability to enter the matrix changes what it does compared to cAMP which only gets into the IMS. Since it is expected that the only differences between the signaling effects of cAMP and 8Br-cAMP relate to IMM permeability and insensitivity to PDEs, it appears that elevated matrix cAMP acts to slow down ATP production, overwhelming any IMS-based upregulation mechanism. These findings are similar to those found by Balaban and co-workers who also used 8Br-cAMP and isolated cardiac mitochondria ^43^, but appear to be at odds with those of Manfredi and co-workers who used 8Br-cAMP and liver mitochondria and concluded that sAC works *only* in the matrix and not the IMS^34^. Manfredi and co-workers also concluded that PKA is the signaling target of sAC and not EPAC as we show here. While it is possible that the discrepancy between our findings and those of others may be cell-type specific, there may be other causes. Importantly, the finding that elevated matrix cAMP (in the form of 8Br-cAMP) slows down ATP production raises important questions about how mitochondrial cAMP microdomains might operate and interact for metabolic regulation.

### Missing link in the mitochondrial sAC signaling cascade

Mechanistically we have tracked the signaling sequence that begins with an elevation of CO_2_/HCO_3_−, followed by AC activation, leading to elevation of cAMP within the IMS, activation of EPAC1 and of Rap1 in the IMS, and eventually to elevation of ATP production by ATP synthase. The localization of these signals is new and important as is their link to augmented ATP production. This raises the next (still unanswered) question: What are the mechanistic details explaining how the Rap1 signal leads to increased ATP production? In particular, does a target of Rap1 carry the message to the ATP synthase directly, or is upregulation of ATP generation achieved indirectly, by acting on some component of the ETC or on a critical membrane transport protein such as ANT or VDAC? Any of these mitochondrial components has the potential to increase ATP production if appropriately activated, but thus far, the relevant mitochondrial targets of Rap1 remain unknown.

### Mitochondrial PKA and CaMKII: location and function

Buck and Levine were the first to suggest that sAC resides in mitochondria, based on experiments in HEK293 cells and a novel antibody for sAC ^32^. Manfredi and co-workers using HeLa, HEK293T, COS-1 cells, and liver mitochondria suggested that sAC was specifically located in the mitochondrial matrix where it activated PKA to affect the electron transport chain ^34^. In contrast, Balaban and co-workers, using pig heart mitochondria presented evidence that PKA was not working at all within any part of cardiac mitochondria to augment ATP production ^43^. In addition, Lefkimmiatis and co-workers showed that PKA does not affect mitochondria matrix signaling but instead that it associates with the cytosolic facing side of the outer mitochondrial membrane ^31, 44^. Here we have presented findings that broadly do not contradict those of Balaban and Lefkimmiatis, but that extend investigation of cAMP signaling in an important new direction. Unlike previous work of others, we included GDP and GTP in our experimental buffers to enable the primary catalytic function of guanine-nucleotide-exchangers such as EPAC as well as small GTPases such as Rap1 ^77^. Additionally, we used the energy substrate pyruvate instead of glutamate. Pyruvate is not only more substantially metabolized by cardiac mitochondria than glutamate but is also metabolized by a larger number of enzymatic steps, which broadens the search for potentially regulated processes. Moreover, we conducted more direct luminescence measurements of mitochondrial ATP production rate, which also provide higher signal-to-noise ratio than the analogous measurements of oxygen consumption presented in previous studies. With this approach we were able to find that cAMP ***does*** act to increase ATP production in cardiac mitochondria but only when it operates in the IMS and acts on EPAC not on PKA.

Another signaling pathway in the mitochondrial matrix was reported by Anderson and his co-workers ^78, 79^. They suggested that mitochondrially localized CaMKII (Ca^2+^/calmodulin-dependent protein kinase II) has multiple targets within mitochondria. Subsequent work by Lezoualc’h and co-authors found that mitochondrial CaMKII was activated by EPAC ^57, 67^. The location of these processes, as presented in these publications, was within the space that includes the intermembrane space (IMS), the inner mitochondrial membrane (IMM) and the matrix, but was not further specified. Our work clearly supports cAMP-dependent activation of EPAC, and it is possible that CaMKII may also be involved. This will be pursued in future work.

### Summary

Here we have demonstrated that there are two independent signaling systems that regulate cardiac mitochondrial ATP production under normal conditions in a complementary manner. One is the well-known calcium signaling pathway that we recently updated ^1, 16, 17^. The second is the newly identified pathway presented here that involves signaling by cAMP, generated in the intermembrane space by soluble adenylyl cyclase (sAC). sAC is activated by bicarbonate, produced from the CO_2_ emission from the consumption of energy substrates. Thus, as substrate consumption increases, a signaling cascade is triggered that results in upregulation of ATP production to meet the increased energy demand. Elements of the cascade have been identified, from sAC to EPAC1 and to Rap1. However, the target(s) of Rap1 in the mitochondrial membranes and/or matrix that cause the increase in ATP production are still unknown. We show that this cAMP signaling pathway complements the calcium-dependent arm of ATP production in the mitochondrial matrix ^1, 16^, and that one or both may be modified in cardiac tissue that has been stressed for prolonged periods following myocardial infarction. It will be important to investigate how each pathway may be modified and contribute to the development of cardiac disease.

## Material and methods

### Mitochondria isolation

Six-to-ten-week-old Sprague-Dawley male rats (250-300 g, from ENVIGO, USA, strain code # 002) were anesthetized using Isoflurane (20 minutes). A thoracotomy and fast excision of the heart was performed, with removal of the atria. The ventricles were minced in ice cold isolation buffer (IB) containing (in mM): KCl 100, MOPS 50, MgSO_4_ 5, EGTA 2, NaPyruvate 10, K_2_HPO_4_ 10. The minced tissue was washed repeatedly with IB until clear of blood. The remainder of the preparation was conducted in a cold room (4°C). 20 mL of IB containing tissue was transferred to a Potter-Elvehjem grinder and homogenized at high speed for 2 seconds followed by 4 repetitive homogenizations with a 1-micron clearance pestle on low speed. The homogenate was centrifuged for 8 min at 600g after which the supernatant was transferred to a new centrifuge tube. The pellet was resuspended with 10 mL IB and centrifuged for 8 min at 600g. The second supernatant was pooled with the first and centrifuged again for 8 min at 600g. The final supernatant was transferred to a clean centrifuge tube and spun at 3200g for 12 min. The supernatant was discarded and the pellet (the mitochondria sample) was resuspended in resuspension buffer (RB1) base solution containing (in mM): KCl 100, MOPS 50, K_2_HPO_4_ 1, supplemented with Na-Pyruvate (10 mM), EGTA (40 μM), and with Fura-2 AM (acetoxymethyl ester form of the calcium indicator Fura-2) (2 μM). After 30 min, the mitochondria were centrifuged at 3200g for 12 min and resuspended in RB2, which is RB supplemented with Na-Pyruvate (1 mM) and EGTA (40 μM). The mitochondria were centrifuged at 3200g for 12 min and a final resuspension and pelleting was done using RB3, consisting of RB and EGTA (40 μM). The concentration of mitochondria was quantified by Lowry assay with a typical rat heart yielding ~15 mg mitochondrial protein. The high purity of mitochondria isolated via this procedure was previously shown ^16,80^. Mitochondria were used within 4 hours of isolation. All procedures and protocols involving animal use were approved by the Institutional Animal Care and Use Committee of the University of Maryland School of Medicine (IACUC # 0921015).

### Isolation of Ventricular Myocytes

Isolated ventricular myocytes were obtained from adult male Sprague-Dawley rats (250-300 g, from ENVIGO, USA, strain code # 002). Animals were euthanized using isoflurane (5%) inhalation anesthesia via a vaporizer. 20 minutes prior to euthanasia rats were injected with an intraperitoneal heparin bolus (1000 U/kg). A thoracotomy was performed, and hearts were excised during deep anesthesia. Hearts were quickly immersed in ice-cold isolation buffer containing 130 mM NaCl, 5.4 mM KCl, 0.5 mM MgCl2, 0.33 mM NaH_2_PO_4_, 10 mM D-glucose, 10 mM taurine, 25 mM HEPES and 0.5 mM EGTA (pH 7.4, adjusted with NaOH). The aorta was cannulated and hearts were then mounted on a Langendorff perfusion system. Hearts were perfused with isolation buffer containing 0.5 mM EGTA buffer for 5 minutes at 37°C, before perfusion was switched to EGTA-free isolation buffer supplemented with 1 mg/mL collagenase (type II; Worthington Biochemical, USA), 0.06 mg/mL protease XIV, 0.06 mg/mL trypsin, and 0.3 mM CaCl_2_. After 6-8 minutes of enzymatic perfusion the heart was dismounted. The ventricles were transferred to isolation buffer supplemented with 2 mg/ml BSA and 20 mM 2,3-butanedione monoxime and cut into small pieces. Mechanical dissociation with a fire polished Pasteur pipette was performed to achieve further dissolution of the ventricular tissue. The cell suspension was then filtered through a nylon mesh with a pore size of 300 μm. Ventricular myocytes were allowed to pellet by sedimentation, resuspended in NT solution, and were used within 4 hours of isolation. All procedures and protocols involving animal use were approved by the Institutional Animal Care and Use Committee of the University of Maryland School of Medicine (IACUC # 0921015).

### Myocardial Infarction Model

Male CD rats (175-200g) underwent surgical ligation of the left anterior descending (LAD) coronary artery, performed by Charles River surgical services. Following incision between the fourth and fifth intercostal spaces, the LAD was permanently ligated between the pulmonary cone and the left auricle using 5-0 silk sutures. As control cohort, age- and size-matched CD rats underwent a sham procedure that included incision but no coronary artery ligation. Post-surgery, transthoracic echocardiography was carried out every two weeks in the MI and sham operated cohorts. All procedures and protocols involving animal use were approved by the Institutional Animal Care and Use Committee of the University of Maryland School of Medicine (IACUC # 0921015).

### Echocardiography

Transthoracic echocardiography was performed using a VisualSonics Vevo 2100. Rats were anesthetized using isoflurane and adjusted to a heart rate of 350 +/− 50 BPM. Body temperature was monitored throughout acquisition. Systolic parameters were obtained from short-axis M-mode scans at the midventricular level, as verified by papillary muscles. Apical four-chamber views were obtained, and diastolic function was assessed by pulsed wave doppler imaging across the mitral valve. B-mode imaging in the parasternal long axis plane was used to calculate global longitudinal strain with VevoStrain software (Visual Sonics). At the end of the acquisition all rats recovered without issues. All procedures and protocols involving animal use were approved by the Institutional Animal Care and Use Committee of the University of Maryland School of Medicine (IACUC # 0921015).

### Immunocytochemistry

200 μL of the ventricular cell suspension (40,000 per mL) were seeded on glass-bottom dishes (Cell Nest, 801001) coated with ECM gel (Sigma, E1270), and allowed to settle for 40 minutes. Cells were fixed with ice-cold methanol for 5 minutes and left to air dry. Cells were then washed with PBS 3 × 5 minutes prior to 2 h of incubation with blocking buffer (SuperBlock™, Thermo #37580). This was followed by overnight incubation(4°C) with primary antibodies against ATP synthase (Abcam, ab128743, 1:50) and sAC (R21 IHC, CEP BIOTECH, 1:50). Cells were then washed 3 × 15 minutes with wash buffer (PBS, 0.2% BSA, 0.05% Triton × (v/v)) and then incubated with appropriate Alexa Fluor™ conjugated secondary antibodies (1:200 in blocking buffer) for 90 minutes at room temperature. Excess antibodies were removed by washing 3 × 15 minutes with wash buffer followed by a 3 × 5 minutes wash with PBS. Cells were then incubated with Alexa Fluor™ 488 Phalloidin (Thermo, A12379, 1:100 in PBS) for 2 h. Excess phalloidin was removed by washing the cells 3 × 15 minutes with PBS.

Isolated cardiac mitochondria were attached to ECM coated glass-bottom 96 well plates by adding 100 μL of 0.01 mg/mL mitochondria to the dish, allowing them to settle for 20 minutes. Mitochondria were fixed for 20 minutes in 100 μL 4% paraformaldehyde in PBS, followed by 3 washes in 200 μL PBS. Membranes were permeabilized utilizing 100 μL 0.05% Triton X-100 in PBS for 15 minutes. Mitochondria were washed 1X with PBS followed by the addition of 100 μL SuperBlock™ solution for 2 hours. Following the removal of blocking buffer, primary antibodies were added in 50 μL blocking buffer for 12 hours at 4° C: ATP synthase antibody (Abcam, ab128743 1:200), sAC antibody (R21 IHC, CEP BIOTECH, 1:50), Tom20 antibody (Santa Cruz, sc-17764, 1:200). After primary antibody incubation, mitochondria were washed 3X with 200 μL PBS. Alexa Fluor™ conjugated secondary antibodies were added in 100 μL blocking buffer at 1:200 for 2 hours at room temperature. Mitochondria were then washed 3X with 200 μL PBS.

### Airy scan sub-diffraction super-resolution imaging

Imaging was carried out with a Zeiss LSM 880 confocal microscope equipped with an Airyscan super resolution imaging module using a 63/1.40 Plan-Apochromat Oil differential interference contrast M27 objective lens (Zeiss MicroImaging). Laser lines at 488-nm (argon laser), 561-nm (diode-pumped solid-state laser) and 633-nm (HeNe laser) were used to detect Alexa Fluor 488, Alexa Fluor 546, and Alexa Fluor 647, respectively. Imaging with each laser line was carried out sequentially. Z stacks of about 0.54 μm depth with intervals of 180 nm were acquired, followed by 3D Airyscan deconvolution to obtain lateral voxel resolution of ~120 nm (at emission centered at 520 nm). Co-localization analysis was done using ZEN image acquisition and processing software (Zeiss MicroImaging; version 2.3SPL). The Pearson’s correlation coefficient (*r*) was used to assess the extent of co-localization between images. Fluorescence intensity profile was analyzed along the transverse axis of ventricular myocytes or the diameter of isolated mitochondria. Fluorescence peak-to-peak distance analysis was carried out using Matlab R2016a.

### Measurements of mitochondrial ATP production and [Ca^2+^]_m_

Measurements of mitochondrial ATP production rate and [Ca^2+^]_m_ were carried out using a BMG LABTECH CLARIOstar plate reader. Fura-2 AM loaded mitochondria (0.1 mg per mL) were mixed in ATP production assay buffer (AB) consisting of (in mM): K-Gluconate 130, KCl 5, K_2_HPO_4_ 1 or 10, MgCl_2_ 1, HEPES 10, EGTA 0.04, GDP 0.125, GTP 0.25, BSA 0.5 mg/mL, D-Luciferin (Sigma) 0.005, Luciferase (Sigma; SRE0045) 0.001 mg/mL, pH 7.2. A luminescence standard curve was performed daily over a range of 100 nM to 1 mM ATP with Oligomycin A (15 μM) treated mitochondria. Where indicated, mitochondria were treated by incubation for 30 minutes prior to the experiment in AB supplemented with the following: [HCO_3_−] (0, 10 or 15 mM, [Na^+^] was kept constant at 15 mM), 1.25 mM [cAMP], 1.25 mM [8br-cAMP], IBMX (0-50 μM), H89 (1 μM), CE3F4 (25 μM). The mitochondria were incubated for 2 minutes prior to the start of the assay with Ca^2+^ (0-50 μM added) and 1 mM Pyruvate and 0.5 mM Malate. Assays were initiated by injection of 100 μL ADP (0.05 or 1.0 mM) and luciferin/luciferase in AB to bring the final volume to 200 μL. Luminescence signal was recorded for 20 seconds with 1 second integration. In the absence of ADP only ~10 nM ATP was present in the system. An automated sequence was used to assess each well first for luminescence then subsequently for fluorescence. ATP production rates were scaled to nMol ATP per sec per mg mitochondrial protein (nMol S^−1^ mg^−1^). [Ca^2+^]_m_ was measured via Fura-2 fluorescence ratio, R_F2_ (excitation: 335 ± 6 nM emission: 490 ± 15 nm / excitation: 380 ± 6 nm, emission: 490 ± 15 nm). The quantitative [Ca^2+^]_m_ values were obtained according to the following equation:

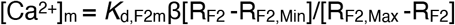

where *K*_d,F2m_ = 0.26 μM, was obtained as described previously ^16^. The β (F_380,min_/ F_380,max_) was measured daily (typically 2.5-2.8), as were Fura-2 R_Max_ (340 nm/380 nm) and R_Min_. Purification of luciferase was critical for accuracy in luminescence measurements for ATP. Briefly, purchased lyophilized powder of luciferase (Sigma, SRE0045) was dissolved to 1 mg/mL in 0.5 M Trizma acetate pH 7.5 (Sigma; T1258). The luciferase in solution was centrifuged at 4° C (14,000g) in filtered centrifugal tubes until the volume within the centrifugal tubes had reduced by 80 % (Amicon Ultra, Millipore, Ireland, molecular cut-off 3 kD). The filtrate was discarded and 0.5 M Trizma acetate solution was added to the luciferase solution at a volume equal to the volume of the discarded filtrate. This wash cycle was repeated 8 times. The concentration of the luciferase stock was re-assessed using a NanoDrop 1000 spectrophotometer (ThermoFisher Scientific), adjusted to 1 mg/mL, and kept at −80° C. Similarly critical was the purification of the ADP stocks as described before ^16^.

### ELISA measurements of cAMP

Quantitative cAMP measurements were carried out on isolated cardiac mitochondria (0.5 mg per sample) using cAMP Direct Immunoassay Kit (Fluorometric) (Abcam, ab138880). Mitochondria samples were treated with 0 or 10 or 15 mM [HCO_3_−], and 0 or 25 μM Forskolin for 30 minutes, pelleted at 3200g for 12 minutes, and lysed using the lysis buffer. Florescence measurements of HRP-cAMP displacement were carried out on the BMG LABTECH CLARIOstar plate reader (excitation: 540 ± 4 nm, emission: 590 ± 4 nm). A standard curve was performed daily over a range of 0.06 pMol to 2 nMol cAMP (0.3 nM to 10 μM cAMP).

### Immune Pull down of Rap1

RalGDS RBD Agarose beads were used to selectively pull-down the active form of Rap1 (Rap-GTP) from isolated cardiac mitochondria (Rap1 Activation Assay Kit, Abcam, ab212011). Samples of mitochondria, 2.5 mg per sample in 2 mL assay buffer, were treated with 0 or 1.25 mM cAMP for 30 minutes on ice, added with Pyruvate (1 mM) and Malate (0.5 mM) and ADP (25 μM), incubated for 2 minutes at room temperature, pelleted by centrifugation at 3200g for 12 minutes, and lysed using the lysis buffer provided with the kit. As positive control, lysed mitochondria samples were treated with 0.1 mM GTPyS (guanosine 5’-O-[gamma-thio]triphosphate) to maximally and irreversibly activate Rap1. Pull down of Rap1-GTP was carried out according to the manufacturer’s instructions. The amounts of active Rap-1 were detected by immunoblot analysis of the pull-down product using anti-Rap1 goat polyclonal antibody (Abcam, ab212011).

### Western blot analysis

Isolated cardiac mitochondria or homogenized freshly harvested ventricular tissue (100–200 μl) were lysed in the same volumes of 2 × SDS sample buffer (Thermo Fisher Scientific) supplemented with 5% β-mercaptoethanol (Millipore) and incubated at 100°C for 10 min, followed by centrifugation at 21,000g at 4°C for 10 min. The supernatants were retained, and protein concentrations were measured directly in these SDS-PAGE samples using a NanoDrop 1000 spectrophotometer (Thermo Fisher Scientific). Proteins were separated on 4–20% gradient Novex Tris-glycine polyacrylamide gels (Thermo Fisher Scientific) and transferred onto polyvinylidene fluoride membranes (Bio-Rad Laboratories). Membranes were blocked in 5% blocking-grade nonfat dry milk (Bio-Rad Laboratories) in PBS-Tween20 and incubated with primary antibodies overnight at 4°C, followed by incubation with secondary antibodies for 60 min at RT. Blots were developed with Super Signal West Pico ECL (Thermo Fisher Scientific) and imaged using Amersham Imager 600 chemiluminescence imager (GE Healthcare Life Sciences). Densitometry quantification was carried out using ImageJ 1.53k (https://imagej.nih.gov/ij/). Primary antibodies used for Western blotting were anti-Tom20 rabbit polyclonal antibody (Proteintech, PTG-11802-AP; 1:20,000), anti Rap1 goat polyclonal antibody (Abcam, ab212011; 1:1,000), OxPhos Human WB Antibody Cocktail for ETC Complexes: CI-20(ND6), C-II-30(FeS), C-III-Core2, C-IV-II and C-V-alpha (ThermoFisher Scientific, 45-8199; 1:2500). Secondary antibodies used were horseradish peroxidase–conjugated anti-mouse (Cell Signaling Technology, CST-7076) or anti-rabbit (Cell Signaling Technology, CST-7074) or donkey anti-goat (Abcam, ab205723).

### Statistics

All results are presented as mean ±SEM. All test conditions are described in detail in the paper. All data collected from experiments in accordance with the specified conditions are included in the paper. All data collected from each experiment was included in the paper. All experiments were repeated independently with at least three separate sample preparations. All the experiments were preformed using sample sizes based on standard protocols in the field. All experiments require fresh heart tissue, thus statistical analysis was carried out in parallel with experiments to determine when further repetition was no longer required. Statistical analysis was performed using either OriginPro 2018 or Matlab R2016a statistical packages, all with α = 0.05. Where appropriate, column analyses were performed using an unpaired, two-tailed t-test (for two groups) or one-way ANOVA with Bonferroni correction (for groups of three or more). Data fitting convergence was achieved with a minimal termination tolerance of 10^−6^. P values less than 0.05 (95% confidence interval) were considered significant. All data displayed a normal distribution and variance was similar between groups for each evaluation.

## Acknowledgments

We thank Joachim Buck, Lonnie Levine and Melania Balbach from Cornell Medical Center for helpful suggestions and their pioneering work on sAC. They have been generous with their resources and insights. We thank Ana Maria Gomez (Université Paris-Saclay & Inserm, UMR-S 1180) for valuable discussions at the beginning of this project. This research was supported by American Heart Association grant 15SDG22100002 (L.B.), 7U19 AI090959 (L.B. & W.J.L.); Frontiers in Anesthesia Research Award from International Anesthesia Research Society (L.B. & W.J.L.); R01 GM129584 (M.K.); University of Maryland Claude D. Pepper Center Grant P30 AG028747 (M.G.); R01 HL142290 (W.J.L); 5R35GM140822 (W.J.L.); U01 HL116321 (W.J.L.); T32 AR007592, the NIH Interdisciplinary Training Grant in Muscle Biology (AKC).

## Material and data availability

We only used commercial materials and resources that are all identified in the Material and methods section (instruments, software, reagents, antibodies, and lab animals). All equations used to analyze the data are identified in the Material and methods section. The data that support the findings of this study are shown within the figures and their source numeric values are included in this publication as supplementary data tables. Should additional information be requested it will be available from the corresponding author.

## Supplementary Information

**Figure S1.**
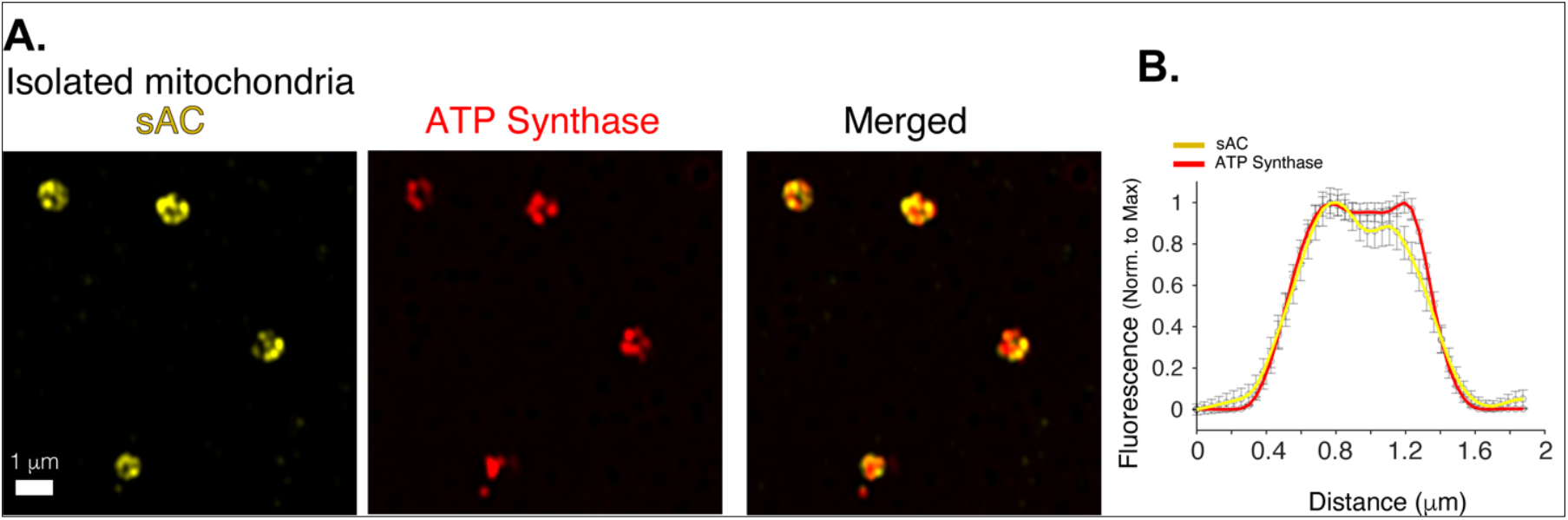
mitochondrial localization of bicarbonate-activated soluble-adenylyl-cyclase (sAC). **A**. AiryScan ‘super-resolution’ images of isolated cardiac mitochondria immuno-labeled for sAC (yellow) and ATP synthase (red). **B**. Immunofluorescence profile of sAC (yellow) and ATP synthase (red) for images as in A, (*n*= 36 mitochondria). Data in **B** are mean ± SEM

**Figure S2.**
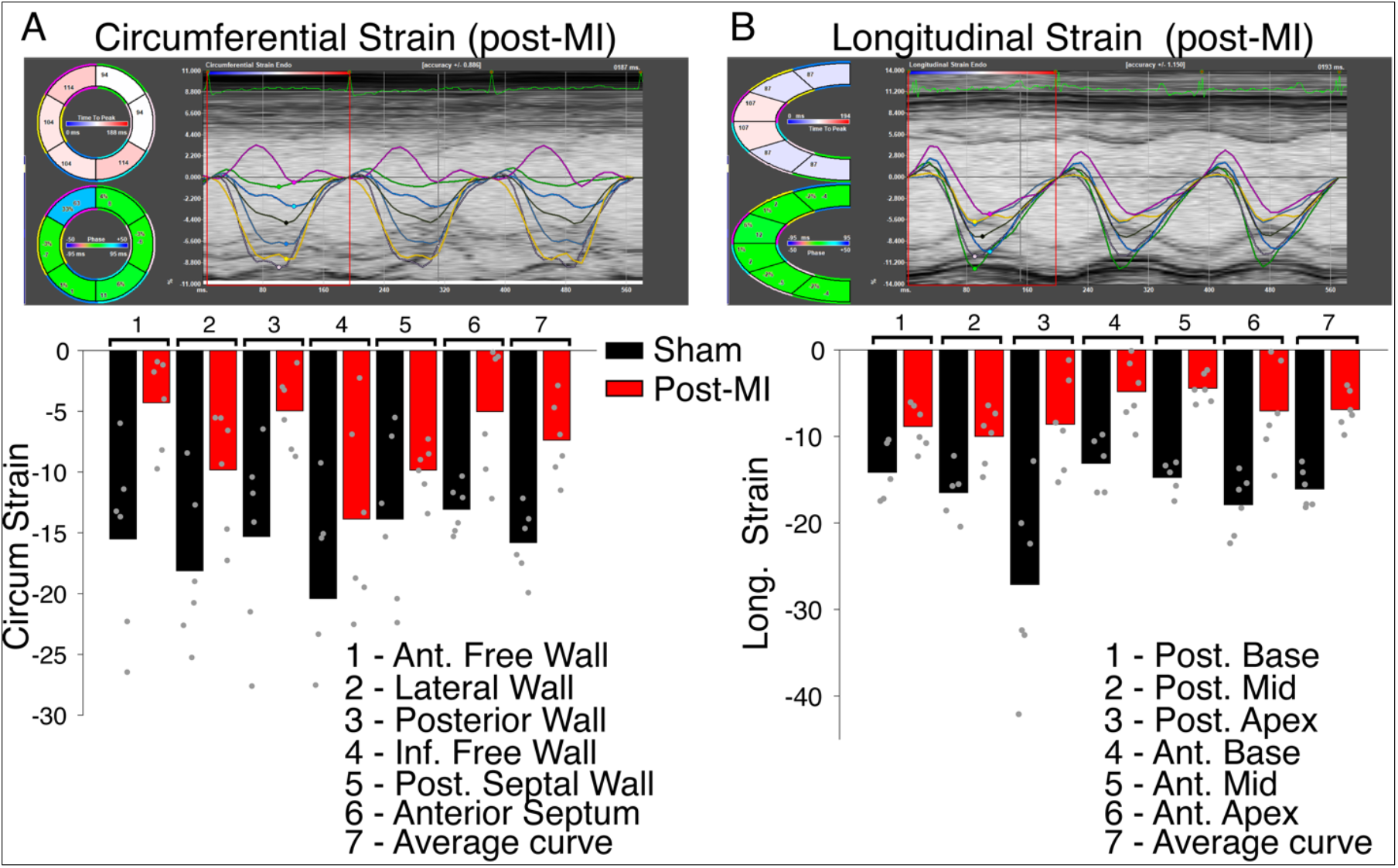
Regional contractile failure detected with speckle-tracking based strain analysis. Time-resolved tracking of myocardial deformation for direct tracking of myocardial strain. **A**. Top panel shows measurements carried out along the circumferential axis 8 weeks post-MI. Six regional strain curves are shown, a seventh **black** line is the average (global) strain at each time point. Bottom panel shows bar charts of the regional and average strain measures (as %) as indicated. Data are from 6 sham operated hearts (black bars) and from 6 hearts with permanent LAD ligation (red bars). Individual data are shown as grey circles. **B**. The same as **A** but measurements taken along the longitudinal axis.

**Figure S3.**
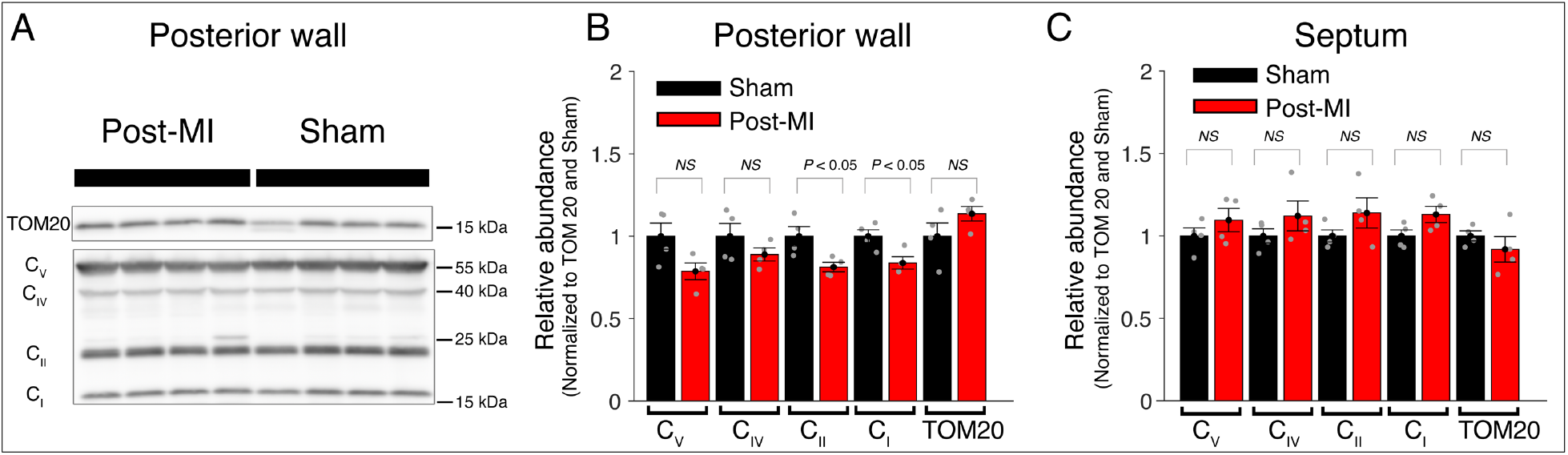
Central components of mitochondrial ATP production are largely unchanged post-MI. **A**. Western blot analysis of mitochondria from the posterior wall 8 weeks after surgery (Post-MI or Sham). Tom20 is used as loading control. C_V_ (ATP synthase subunit 5A), C_IV_ (complex 4 MTCO1), C_II_ (complex 2 SDHB), C_I_ (complex 1 NDUFB8). **B**. Relative abundance of respiratory chain proteins in mitochondria isolated from the posterior wall 8 weeks after surgery (Red bars = Post-MI, Black bars = Sham). **C**. Same as B but mitochondria are isolated from the septum. Data are mean ± SEM. Two-tailed t-test. Figure S3—source data 1. Western blot analysis in Figure S3 (anti-Cv, anti-Civ, anti-Cii, anti-Ci, anti-TOM20). Figure S3—source data 2. Original file for western blot analysis of posterior wall in Figure S3 (anti-Cv, anti-Civ, anti-Cii, anti-Ci). Figure S3—source data 3. Original file for western blot analysis of posterior wall in Figure S3 (anti-TOM20). Figure S3—source data 4. Original file for western blot analysis of septal wall in Figure S3 (anti-Cv, anti-Civ, anti-Cii, anti-Ci). Figure S3—source data 5. Original file for western blot analysis of septal wall in Figure S3 (anti-TOM20).

**Supplementary table 1.** Mechanistic findings by investigations of sAC and its role in mitochondria.

| Citation | Native Mitochondrial sAC localization | Role of PKA | Physiological role of sAC signaling |
| --- | --- | --- | --- |
| **Manfredi** and coworkers<sup>34, 40-42</sup> | Matrix | PKA-dependent | Sensing nutrient availability |
| **Balaban** and coworkers<sup>43</sup> | ND | PKA-independent | ND |
| **Lefkimmatis**<br>23, 44 | ND | PKA-independent | ND |
| **Brenner** and coworkers<sup>45</sup> | ND | PKA-independent | Regulates calcium accumulation, permeability transition and cell death |
| This paper | mitochondrial inter membrane space | PKA-independent | Sensing substrate consumption - mechano-metabolic sensor |

## References

1. Boyman, L., Karbowski, M. & Lederer, W.J. Regulation of Mitochondrial ATP Production: Ca(2+) Signaling and Quality Control. Trends Mol Med 26, 21–39 (2020).

2. From, A.H., Petein, M.A., Michurski, S.P., Zimmer, S.D. & Ugurbil, K. 31P-NMR studies of respiratory regulation in the intact myocardium. FEBS Lett 206, 257–261 (1986).

3. From, A.H. et al. Regulation of the oxidative phosphorylation rate in the intact cell. Biochemistry 29, 3731–3743 (1990).

4. Katz, L.A., Swain, J.A., Portman, M.A. & Balaban, R.S. Relation between phosphate metabolites and oxygen consumption of heart in vivo. Am J Physiol 256, H265–274 (1989).

5. Portman, M.A., Heineman, F.W. & Balaban, R.S. Developmental changes in the relation between phosphate metabolites and oxygen consumption in the sheep heart in vivo. J Clin Invest 83, 456–464 (1989).

6. Zhang, J. et al. Transmural bioenergetic responses of normal myocardium to high workstates. Am J Physiol 268, H1891–1905 (1995).

7. Elliott, A.C., Smith, G.L. & Allen, D.G. The metabolic consequences of an increase in the frequency of stimulation in isolated ferret hearts. J Physiol 474, 147–159 (1994).

8. Matthews, P.M., Bland, J.L., Gadian, D.G. & Radda, G.K. The steady-state rate of ATP synthesis in the perfused rat heart measured by 31P NMR saturation transfer. Biochem Biophys Res Commun 103, 1052–1059 (1981).

9. Massie, B.M. et al. Myocardial metabolism during increased work states in the porcine left ventricle in vivo. Circ Res 74, 64–73 (1994).

10. Schwartz, G.G., Greyson, C.R., Wisneski, J.A., Garcia, J. & Steinman, S. Relation among regional O2 consumption, high-energy phosphates, and substrate uptake in porcine right ventricle. Am J Physiol 266, H521–530 (1994).

11. Xu, Y., Lu, L., Zhu, P. & Schwartz, G.G. beta-adrenergic stimulation induces transient imbalance between myocardial substrate uptake and metabolism in vivo. Am J Physiol 275, H2181–2190 (1998).

12. Jacobus, W.E. Respiratory control and the integration of heart high-energy phosphate metabolism by mitochondrial creatine kinase. Annu Rev Physiol 47, 707–725 (1985).

13. Wang, Z. et al. Specific metabolic rates of major organs and tissues across adulthood: evaluation by mechanistic model of resting energy expenditure. Am J Clin Nutr 92, 1369–1377 (2010).

14. Maack, C. & O’Rourke, B. Excitation-contraction coupling and mitochondrial energetics. Basic Res Cardiol 102, 369–392 (2007).

15. Murphy, E. & Steenbergen, C. Regulation of Mitochondrial Ca(2+) Uptake. Annu Rev Physiol 83, 107–126 (2021).

16. Wescott, A.P., Kao, J.P.Y., Lederer, W.J. & Boyman, L. Voltage-energized Calcium-sensitive ATP Production by Mitochondria. Nat Metab 1, 975–984 (2019).

17. Garg, V. et al. The mechanism of MICU-dependent gating of the mitochondrial Ca(2+)uniporter. eLife 10(2021).

18. Zhao, G., Joca, H.C., Nelson, M.T. & Lederer, W.J. ATP- and voltage-dependent electro-metabolic signaling regulates blood flow in heart. Proc Natl Acad Sci U S A 117, 7461–7470 (2020).

19. Grainger, N. & Santana, L.F. Metabolic-electrical control of coronary blood flow. Proc Natl Acad Sci U S A 117, 8231–8233 (2020).

20. Buck, J., Sinclair, M.L., Schapal, L., Cann, M.J. & Levin, L.R. Cytosolic adenylyl cyclase defines a unique signaling molecule in mammals. Proc Natl Acad Sci U S A 96, 79–84 (1999).

21. Litvin, T.N., Kamenetsky, M., Zarifyan, A., Buck, J. & Levin, L.R. Kinetic properties of “soluble” adenylyl cyclase. Synergism between calcium and bicarbonate. J Biol Chem 278, 15922–15926 (2003).

22. Zaccolo, M., Filippin, L., Magalhaes, P. & Pozzan, T. Heterogeneity of second messenger levels in living cells. Novartis Found Symp 239, 85–93; discussion 93–85, 150–159 (2001).

23. Di Benedetto, G., Lefkimmiatis, K. & Pozzan, T. The basics of mitochondrial cAMP signalling: Where, when, why. Cell Calcium 93, 102320 (2021).

24. Oldham, W.M. & Hamm, H.E. Heterotrimeric G protein activation by G-protein-coupled receptors. Nat Rev Mol Cell Biol 9, 60–71 (2008).

25. Syrovatkina, V., Alegre, K.O., Dey, R. & Huang, X.Y. Regulation, Signaling, and Physiological Functions of G-Proteins. J Mol Biol 428, 3850–3868 (2016).

26. Scott, J.D., Dessauer, C.W. & Tasken, K. Creating order from chaos: cellular regulation by kinase anchoring. Annu Rev Pharmacol Toxicol 53, 187–210 (2013).

27. Zaccolo, M. & Pozzan, T. Discrete microdomains with high concentration of cAMP in stimulated rat neonatal cardiac myocytes. Science 295, 1711–1715 (2002).

28. Lomas, O. & Zaccolo, M. Phosphodiesterases maintain signaling fidelity via compartmentalization of cyclic nucleotides. Physiology (Bethesda) 29, 141–149 (2014).

29. Baillie, G.S., Tejeda, G.S. & Kelly, M.P. Therapeutic targeting of 3’,5’-cyclic nucleotide phosphodiesterases: inhibition and beyond. Nat Rev Drug Discov 18, 770–796 (2019).

30. Mongillo, M. et al. Fluorescence resonance energy transfer-based analysis of cAMP dynamics in live neonatal rat cardiac myocytes reveals distinct functions of compartmentalized phosphodiesterases. Circ Res 95, 67–75 (2004).

31. Burdyga, A. et al. Phosphatases control PKA-dependent functional microdomains at the outer mitochondrial membrane. Proc Natl Acad Sci U S A 115, E6497–E6506 (2018).

32. Zippin, J.H. et al. Compartmentalization of bicarbonate-sensitive adenylyl cyclase in distinct signaling microdomains. FASEB J 17, 82–84 (2003).

33. Feng, Q. et al. Two domains are critical for the nuclear localization of soluble adenylyl cyclase. Biochimie 88, 319–328 (2006).

34. Acin-Perez, R. et al. Cyclic AMP produced inside mitochondria regulates oxidative phosphorylation. Cell Metab 9, 265–276 (2009).

35. Tresguerres, M., Levin, L.R. & Buck, J. Intracellular cAMP signaling by soluble adenylyl cyclase. Kidney Int 79, 1277–1288 (2011).

36. Rossetti, T., Jackvony, S., Buck, J. & Levin, L.R. Bicarbonate, carbon dioxide and pH sensing via mammalian bicarbonate-regulated soluble adenylyl cyclase. Interface focus 11, 20200034 (2021).

37. Schmid, A. et al. Soluble adenylyl cyclase is localized to cilia and contributes to ciliary beat frequency regulation via production of cAMP. J Gen Physiol 130, 99–109 (2007).

38. Sample, V. et al. Regulation of nuclear PKA revealed by spatiotemporal manipulation of cyclic AMP. Nat Chem Biol 8, 375–382 (2012).

39. Parker, T., Wang, K.W., Manning, D. & Dart, C. Soluble adenylyl cyclase links Ca(2+) entry to Ca(2+)/cAMP-response element binding protein (CREB) activation in vascular smooth muscle. Scientific reports 9, 7317 (2019).

40. Valsecchi, F., Ramos-Espiritu, L.S., Buck, J., Levin, L.R. & Manfredi, G. cAMP and mitochondria. Physiology (Bethesda) 28, 199–209 (2013).

41. Valsecchi, F., Konrad, C. & Manfredi, G. Role of soluble adenylyl cyclase in mitochondria. Biochim Biophys Acta 1842, 2555–2560 (2014).

42. Acin-Perez, R. et al. A phosphodiesterase 2A isoform localized to mitochondria regulates respiration. J Biol Chem 286, 30423–30432 (2011).

43. Covian, R., French, S., Kusnetz, H. & Balaban, R.S. Stimulation of oxidative phosphorylation by calcium in cardiac mitochondria is not influenced by cAMP and PKA activity. Biochim Biophys Acta 1837, 1913–1921 (2014).

44. Lefkimmiatis, K., Leronni, D. & Hofer, A.M. The inner and outer compartments of mitochondria are sites of distinct cAMP/PKA signaling dynamics. J Cell Biol 202, 453–462 (2013).

45. Wang, Z. et al. A cardiac mitochondrial cAMP signaling pathway regulates calcium accumulation, permeability transition and cell death. Cell Death Dis 7, e2198 (2016).

46. Di Benedetto, G., Scalzotto, E., Mongillo, M. & Pozzan, T. Mitochondrial Ca(2)(+) uptake induces cyclic AMP generation in the matrix and modulates organelle ATP levels. Cell Metab 17, 965–975 (2013).

47. van Dyke, R.W., Gollan, J.L. & Scharschmidt, B.F. Oxygen consumption by rat liver: effects of taurocholate and sulfobromophthalein transport, glucagon, and cation substitution. Am J Physiol 244, G523–531 (1983).

48. Bracht, L., Caparroz-Assef, S.M., Bracht, A. & Bersani-Amado, C.A. Effect of the Combination of Ezetimibe and Simvastatin on Gluconeogenesis and Oxygen Consumption in the Rat Liver. Basic Clin Pharmacol Toxicol 118, 415–420 (2016).

49. Shadrin, K.V., Morgulis, II, Pahomova, V.G., Rupenko, A.P. & Khlebopros, R.G. Characteristics of oxygen transport through the surface of the isolated perfused rat liver. Dokl Biochem Biophys 464, 298–300 (2015).

50. do Nascimento, G.S. et al. The acute effects of citrus flavanones on the metabolism of glycogen and monosaccharides in the isolated perfused rat liver. Toxicol Lett 291, 158–172 (2018).

51. de Medeiros, H.C., Constantin, J., Ishii-Iwamoto, E.L. & Mingatto, F.E. Effect of fipronil on energy metabolism in the perfused rat liver. Toxicol Lett 236, 34–42 (2015).

52. Colturato, C.P. et al. Metabolic effects of silibinin in the rat liver. Chem Biol Interact 195, 119–132 (2012).

53. Rostovtseva, T. & Colombini, M. ATP flux is controlled by a voltage-gated channel from the mitochondrial outer membrane. J Biol Chem 271, 28006–28008 (1996).

54. Rostovtseva, T. & Colombini, M. VDAC channels mediate and gate the flow of ATP: implications for the regulation of mitochondrial function. Biophys J 72, 1954–1962 (1997).

55. Rostovtseva, T.K., Komarov, A., Bezrukov, S.M. & Colombini, M. Dynamics of nucleotides in VDAC channels: structure-specific noise generation. Biophys J 82, 193–205 (2002).

56. Zippin, J.H. et al. CO2/HCO3(−)- and calcium-regulated soluble adenylyl cyclase as a physiological ATP sensor. J Biol Chem 288, 33283–33291 (2013).

57. Fazal, L. et al. Multifunctional Mitochondrial Epac1 Controls Myocardial Cell Death. Circ Res 120, 645–657 (2017).

58. Liu, D. et al. PDE2 regulates membrane potential, respiration and permeability transition of rodent subsarcolemmal cardiac mitochondria. Mitochondrion 47, 64–75 (2019).

59. Chen, J., Martinez, J., Milner, T.A., Buck, J. & Levin, L.R. Neuronal expression of soluble adenylyl cyclase in the mammalian brain. Brain Res 1518, 1–8 (2013).

60. Barth, E., Stammler, G., Speiser, B. & Schaper, J. Ultrastructural quantitation of mitochondria and myofilaments in cardiac muscle from 10 different animal species including man. J Mol Cell Cardiol 24, 669–681 (1992).

61. Omura, T. Mitochondria-targeting sequence, a multi-role sorting sequence recognized at all steps of protein import into mitochondria. J Biochem 123, 1010–1016 (1998).

62. Zhang, G., Liu, Y., Ruoho, A.E. & Hurley, J.H. Structure of the adenylyl cyclase catalytic core. Nature 386, 247–253 (1997).

63. Wells, J.N. & Miller, J.R. Methylxanthine inhibitors of phosphodiesterases. Methods Enzymol 159, 489–496 (1988).

64. Maurice, D.H. et al. Advances in targeting cyclic nucleotide phosphodiesterases. Nat Rev Drug Discov 13, 290–314 (2014).

65. Valsecchi, F. et al. Distinct intracellular sAC-cAMP domains regulate ER Ca(2+) signaling and OXPHOS function. J Cell Sci 130, 3713–3727 (2017).

66. Kumar, S., Kostin, S., Flacke, J.P., Reusch, H.P. & Ladilov, Y. Soluble adenylyl cyclase controls mitochondria-dependent apoptosis in coronary endothelial cells. J Biol Chem 284, 14760–14768 (2009).

67. Laudette, M. et al. Cyclic AMP-binding protein Epac1 acts as a metabolic sensor to promote cardiomyocyte lipotoxicity. Cell Death Dis 12, 824 (2021).

68. Christensen, A.E. et al. cAMP analog mapping of Epac1 and cAMP kinase. Discriminating analogs demonstrate that Epac and cAMP kinase act synergistically to promote PC-12 cell neurite extension. J Biol Chem 278, 35394–35402 (2003).

69. Kawasaki, H. et al. A family of cAMP-binding proteins that directly activate Rap1. Science 282, 2275–2279 (1998).

70. de Rooij, J. et al. Epac is a Rap1 guanine-nucleotide-exchange factor directly activated by cyclic AMP. Nature 396, 474–477 (1998).

71. Luongo, T.S. et al. The Mitochondrial Calcium Uniporter Matches Energetic Supply with Cardiac Workload during Stress and Modulates Permeability Transition. Cell reports 12, 23–34 (2015).

72. Kwong, J.Q. et al. The Mitochondrial Calcium Uniporter Selectively Matches Metabolic Output to Acute Contractile Stress in the Heart. Cell reports 12, 15–22 (2015).

73. Liu, T., Yang, N., Sidor, A. & O’Rourke, B. MCU Overexpression Rescues Inotropy and Reverses Heart Failure by Reducing SR Ca(2+) Leak. Circ Res 128, 1191–1204 (2021).

74. Pan, X. et al. The physiological role of mitochondrial calcium revealed by mice lacking the mitochondrial calcium uniporter. Nat Cell Biol 15, 1464–1472 (2013).

75. Gómez, A.M., Guatimosim, S., Dilly, K.W., Vassort, G. & Lederer, W.J. Heart failure after myocardial infarction: altered excitation-contraction coupling. Circulation 104, 688–693 (2001).

76. Alvarez, J.L., Aimond, F., Lorente, P. & Vassort, G. Late post-myocardial infarction induces a tetrodotoxin-resistant Na(+)Current in rat cardiomyocytes. J Mol Cell Cardiol 32, 1169–1179 (2000).

77. Traut, T.W. Physiological concentrations of purines and pyrimidines. Mol Cell Biochem 140, 1–22 (1994).

78. Joiner, M.L. et al. CaMKII determines mitochondrial stress responses in heart. Nature 491, 269–273 (2012).

79. Luczak, E.D. et al. Mitochondrial CaMKII causes adverse metabolic reprogramming and dilated cardiomyopathy. Nat Commun 11, 4416 (2020).

80. Shimada, B.K. et al. Pyruvate-Driven Oxidative Phosphorylation is Downregulated in Sepsis-Induced Cardiomyopathy: A Study of Mitochondrial Proteome. Shock 57, 553–564 (2022).

